# Promoting extinction or minimizing growth? The impact of treatment on trait trajectories in evolving populations

**DOI:** 10.1101/2022.06.17.496570

**Authors:** Michael Raatz, Arne Traulsen

## Abstract

When cancers or bacterial infections establish, small populations of cells have to free themselves from homoeostatic regulations that prevent their expansion. Trait evolution allows these populations to evade this regulation, escape stochastic extinction and climb up the fitness landscape. In this study, we analyse this complex process and investigate the fate of a cell population that underlies the basic processes of birth, death and mutation. We find that the shape of the fitness landscape dictates a circular adaptation trajectory in the trait space spanned by birth and death rates. We show that successful adaptation is less likely for parental populations with higher turnover (higher birth and death rates). Including density- or trait-affecting treatment we find that these treatment types change the adaptation dynamics in agreement with a geometrical analysis of fitness gradients. Treatment strategies that simultaneously target birth and death rates are most effective, but also increase evolvability. By mapping physiological adaptation pathways and molecular drug mechanisms to traits and treatments with clear eco-evolutionary consequences, we can achieve a much better understanding of the adaptation dynamics and the eco-evolutionary mechanisms at play in the dynamics of cancer and bacterial infections.

## 1. Introduction

Cancer cells and bacterial pathogens show extensive adaptive potential, which helps them to establish even in unfavourable conditions and outgrow competitors and external pressures, for example by the immune system (Fridman et al., 2012; Winstanley et al., 2016). In healthy tissue or healthy microbiomes, external regulation aims to maintain a constant population size, which together with stochastic fluctuations in the population dynamics of individual subpopulations results in a constant turnover characterized by the eventual stochastic extinction of a specific subpopulation and subsequent replacement by other subpopulations (Gallaher et al., 2019). This extinction can be prevented by adaptations that give an emerging subpopulation of cells a fitness advantage over the remaining population. The increased fitness reduces the subpopulation’s risk of extinction in a process often termed evolutionary rescue (Orr and Unckless, 2008; Alexander et al., 2014; Uecker et al., 2014; Marrec and Bitbol, 2020a). Accordingly, the onset of cancer is characterized by malignant cells breaking with the homoeostatic regulation of healthy tissue (Basanta and Anderson, 2013, 2017). Similarly, bacterial infections that either emerge from or invade an otherwise healthy microbiome have to develop mechanisms to outgrow the other community members and free themselves from regulative community interactions, for example by pathoadaptive mutations (Winstanley et al., 2016; Culyba and Tyne, 2021).

Many individual mechanisms of how this fitness increase is realized have been identified. In a progressing tumour, the net growth increase of subclones relative to their parental clones often indicates a continuing evolution towards higher net growth rates, often but not always driven by the accumulation of known driver mutations (Gruber et al., 2019). Biswas et al. (2004) suggest that NF-*κ*B activation increases proliferation and decreases apoptosis rate in estrogen receptor-negative breast cancer cells. Lopez and Tait (2015) describe how apoptosis is avoided in cancer cells by upregulating anti-apoptotic BCL-2 proteins. Similary, also infectious bacteria must adapt during an ongoing infection (Faure et al., 2018; Culyba and Tyne, 2021). For example, Young et al. (2017) showed that formerly commensal constituents of the host microbiome accrue substantial adaptive genotypic changes as they become infective, and Both et al. (2021) documented the phenotypic changes during the adaptation to the host environment.

These adaptations have led to the development of drugs that target many such mechanisms both in cancer and in bacterial infections. For example, BCL-2 inhibitors aim to counter decreased apoptosis rates in cancer cells (Montero and Letai, 2018), and NF-*κ*B inhibition is investigated to lessen the inflammatory increase in proliferation (Yu et al., 2020). Anti-virulence therapy and microbiome modulation have been proposed as options besides antibiotics to counter the adaptations of pathogenic bacteria (Hauser et al., 2016).

The diversity of these specific, experimentally well-characterized adaptations and potential treatments call for an abstraction to elucidate the eco-evolutionary mechanisms behind adaptations of cell populations in challenging environments. It is a priori unclear which functional traits of cancer cells or pathogenic bacteria would be targeted by adaptations. Similarly, it is not understood how treatment-induced perturbations to the adapting populations or their environments would affect the adaptation process. In order to generalize from the plethora of adaptive mutations or plastic responses of cancer cells and bacterial pathogens, we describe the population of evolving cells in a minimal model: Cells competitively grow, die and mutate. We speculate that many of the adaptive mechanisms described above can be classified as either increasing the birth rate or decreasing the death rate. Treatment approaches that try to contain or eradicate such adapting populations could then be grouped into two types: (i) They either directly decrease the population size of the target population, or (ii) indirectly decrease the population size by affecting their birth and death rates. In such a simplistic but general setting we investigate where adaptation will take the population in a trait space spanned by birth rate and death rate, and how treatment will affect the resulting adaptation trajectories.

## 2. Methods

### 2.1. Description of the underlying microscopic processes

We represent the initial phases of tumour formation or the establishment of a bacterial infection as the spread of a population of cells in a harsh environment. In our model, this harshness manifests in similar birth and death rates and a decreasing birth rate as population size increases. The similarity of birth and death rates is supported by the high proportion of dead cells in tumours (Kerr and Lamb, 1984; de Jong et al., 2000; Liu et al., 2001; Alenzi, 2004; Gallaher et al., 2019). While bacterial death rates in benign conditions are small (Koch, 1959; Stewart et al., 2005) the mortality from immune responses or nutrient scarcity may be considerable and the importance of bacterial death is probably underestimated (Frenoy and Bonhoeffer, 2018). Oxygen availability, space restriction and nutrient limitation are likely mechanisms for the density dependence of the birth rate. We assume that this density dependence restricts the birth rate *β* of cells by a logistic term with a carrying capacity *K*. We assume that death occurs at a constant rate *δ*. Upon each birth event mutations can give rise to lineages with trait combinations (*β*_*m*_, *δ*_*m*_) that slightly deviate from those of their parental lineage. We assume that mutations in the two traits can occur independently and without correlation, and that mutational effects are purely additive. The birth and expansion of fitter mutants can shift the population average trait combination and thus cause the population to adapt by exploring its adaptive landscape (e.g. Patout et al., 2021). We can represent the adaptation of a population by the trajectory of the mean trait combination in the trait space spanned by birth rate *β* and death rate *δ*. Focussing on the initial phases of adaptation, we assume that the carrying capacity *K* remains constant. We will investigate treatment types that either target the density or the traits of the evolving population (Fig. 1). Density-affecting treatment types are modelled as instantaneous density reductions (bottlenecks) applied homogeneously to the whole population, similar to the resection of a tumour where cancerous tissue is surgically removed, or the voiding of the bladder during urinary tract infections where most non-attached pathogenic bacteria are flushed out (Cox and Hinman, 1961; Sobel, 1997). Trait-affecting treatment types are implemented by prolonged additive changes to either the birth or the death rates of the individual lineages. ‘Static’ drugs decrease the birth rate by Δ_*β*_ (e.g. cytostatic chemotherapy or bacteriostatic antibiotics), ‘toxic’ drugs increase the death rate byΔ_*δ*_ (e.g. cytotoxic chemotherapy, immunotherapy or bactericidal antibiotics). Different trait-affecting treatment types can thus be represented by vectors (Δ_*β*_, Δ_*δ*_) in trait space (Fig. 1). Accounting for treatment and logistic density dependence of birth rates the effective birth and death rates of lineage *i* with population size *N*_*i*_ are given by

**Figure 1.**
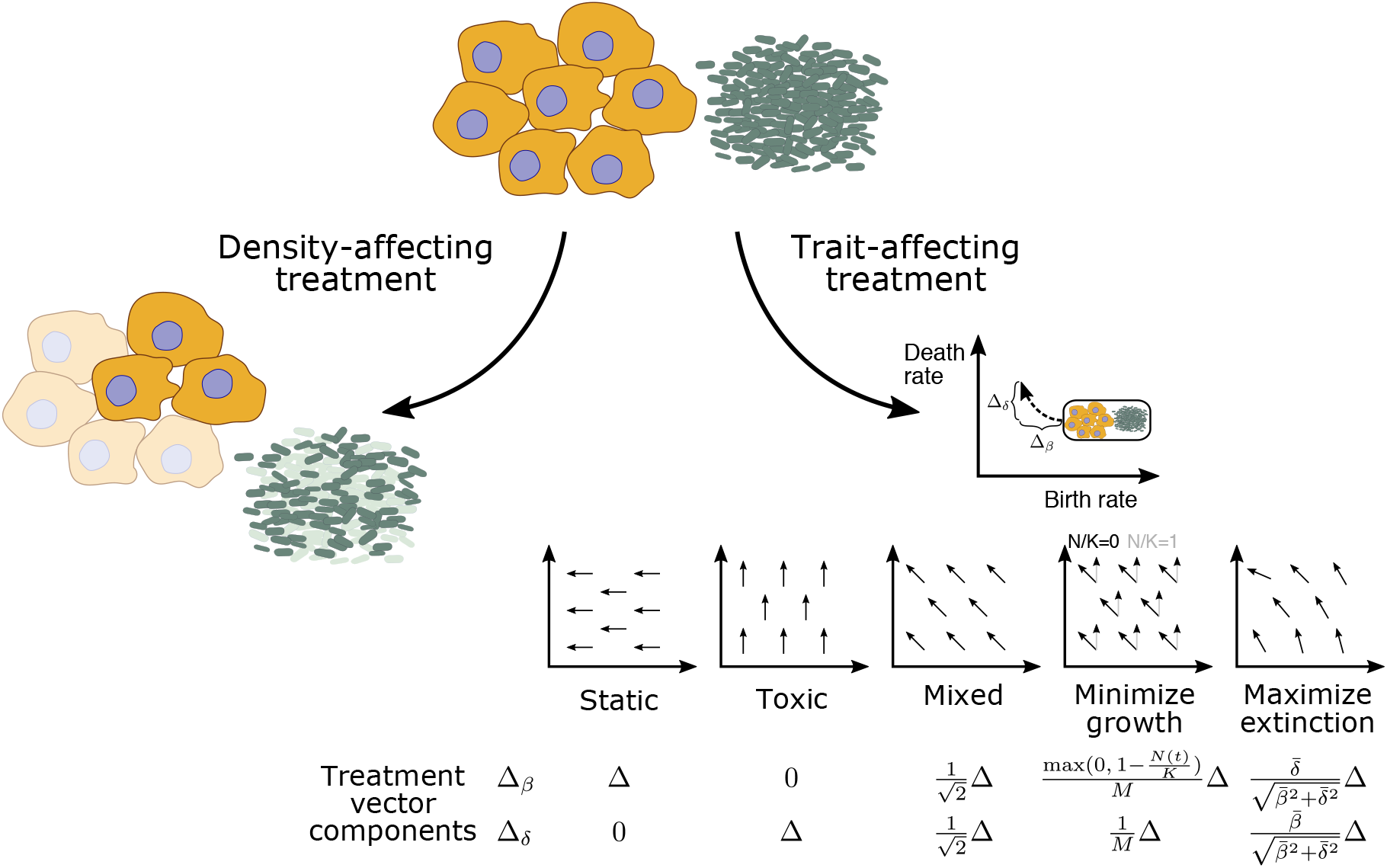
Different treatment types can either affect the cell density directly (left) or indirectly via changing the traits (right). Populations of cancer cells (yellow) or pathogenic bacteria (green) can be targeted with different mechanisms. Density-affecting treatment applies a bottleneck and reduces the population size instantaneously to a fraction *f*. Trait-affecting treatment, e.g. chemotherapy, alters the traits for a prolonged time period (the treatment duration) and displaces the population in trait space temporarily which results in population decline. Note that 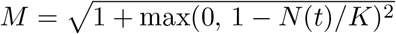 is a normalization factor.

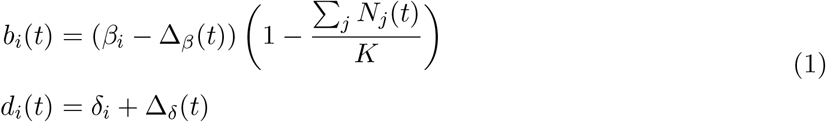

We ensure that effective birth rates are always greater than or equal to zero, setting them to zero if they would be negative.

### 2.2. Stochastic model

We use these microprocesses of birth, death and mutation to construct a discrete-time stochastic model (Eq. 2). We assume that the number of birth and death events per lineage *i* per time step *dt*, (*B*_*i*_(*t*+*dt*) and *D*_*i*_(*t* + *dt*)) are Poisson-distributed around the expected numbers of birth events *b*_*i*_ *N*_*i*_ *dt* and death events *d*_*i*_ *N*_*i*_ *dt*, given the effective birth and death rates *b*_*i*_ and *d*_*i*_ according to Eq. (1). The number of mutants *M*_*i*_(*t* + *dt*) among the new-born cells is given by a binomial distribution with mutation probability *μ*.

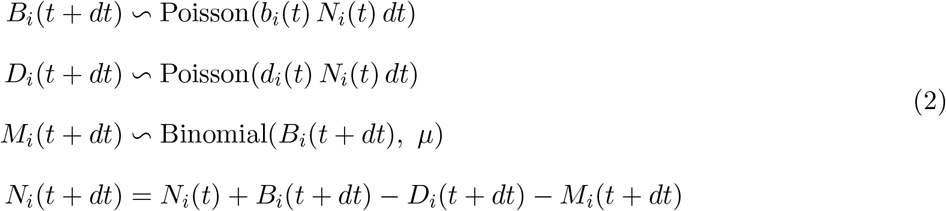

Each newly mutated cell founds a new lineage with trait values drawn from a truncated Gaussian distribution with the parental trait values as the mean and a standard deviation of *σ* = 0.05. By setting the lower bound of the truncated Gaussian distribution to zero, we prevent the evolution of negative trait values. The upper bound was set to 1000, which is far beyond the trait values that are obtained in our simulations and thus does not affect our results. By assuming a truncated Gaussian distribution of mutational effects we draw the mutant trait values predominantly from the vicinity of the parental traits. Thus, we focus our investigation on an adaptive process where trait changes occur predominantly in small steps, either by plastic changes to the cell phenotypes or by mutations with small effects, albeit single large-effect jackpot events are also possible but much less likely. This represents the diversity of adaptive mechanisms in cancer and pathoadaptations in bacterial infections resulting from the multitude of stressors that adapting cell populations face in the human body.

### 2.3. Deterministic model

Defining the total population size as *N* (*t*) =∑ _*i*_ *N*_*i*_(*t*) and the population average traits as 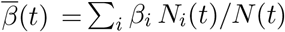 and 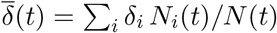, we can construct a deterministic model from the above microscopic model using a Quantitative Genetics approach (Lande, 1982),

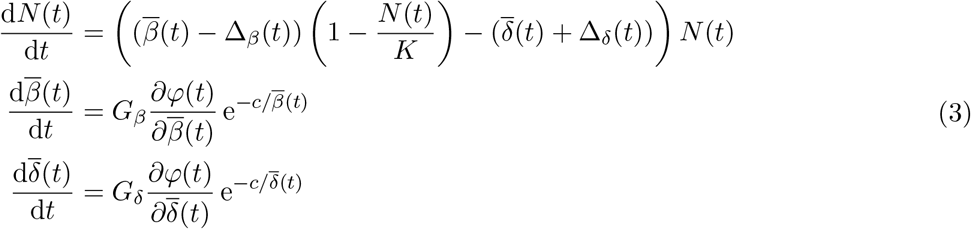

Here, the change in total population size is governed by the difference of logistic average birth rate and average death rate. Treatment affects the effective birth and death rates as in Eq. (1). The change in the average birth and death rates are assumed to be proportional to the gradient of a function *φ*(*t*) (defined below) that describes the fitness of individuals with proportionality constants *G*_*β*_ and *G*_*δ*_ that describe the additive genetic variance in the traits (Lande, 1982). The factors 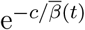 and 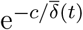 ensure decelerating trait changes close to the trait axis, thus preventing negative trait values (Abrams and Matsuda, 1997; Raatz et al., 2019). Note that also this deterministic model formulation assumes independence of the two traits. The system of ordinary differential equations Eq. 3 is numerically integrated using the LSODA implementation of the solve ivp function from the Scipy library (Virtanen et al., 2020) in Python (version 3.8). Standard initial conditions are *N* (0) = 100, 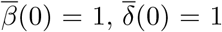 (Tab. 1).

**Table 1.** Reference parameter set. The parameters of the stochastic adaptive process are chosen such that without treatment about half of the replicate simulations show successful adaptation. The parameters of the deterministic model were set such that the time scales of the deterministic dynamics would match the time scales of the stochastic model. Deviations from these values are reported where applicable.

| Parameter | Biological meaning | Value |
| --- | --- | --- |
| $\beta_0$ | Birth rate of the first parental lineage | 1 |
| $\delta_0$ | Death rate of the first parental lineage | 1 |
| $K$ | Carrying capacity for total cell population | 20 000 |
| $\Delta$ | Absolute treatment effect in trait space | 0.5 |
| $N_0$ | Initial size of the first parental lineage | 100 |
| $dt$ | Time step for evaluation of stochastic dynamics | 0.1 |
| $\mu$ | Mutation probability per cell division | 0.005 |
| $\sigma$ | Standard deviation of mutational trait changes | 0.05 |
| $G_\beta$ | Genetic variance of birth rate | $10^{-2.5}$ |
| $G_\delta$ | Genetic variance of death rate | $10^{-2.5}$ |
| $c$ | Trait change deceleration for small trait values | 0.1 |

Setting the temporal derivative of the population size to zero we can obtain the conditions for the manifold where the population change equals zero. On this manifold, the population size is given by the effective carrying capacity

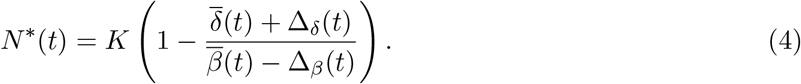

Because of treatment, the effective carrying capacity could become negative. In our simulations, however, we ensure that the population size remains bounded by zero.

### 2.4. Defining fitness

Adaptation should increase fitness relative to competitors, but what exactly determines fitness in populations that have to adapt to unfavourable conditions? Generally, defining fitness measures is ambiguous (Doebeli et al., 2017; Kokko, 2021). One possible definition is lifetime-reproductive output, which itself is a composite measure that includes net growth rate, but also the probability that newly founded lineages survive stochastic population size fluctuations. Even in our simplified setting the determinants of fitness are a priori not trivial, particularly in a regime of high rates of stochastic extinction of lineages. An obvious choice may be the net growth of a lineage *r*, which determines how quickly that lineage grows out of this regime of probable stochastic extinction and outcompetes other lineages. Similarly, the survival probability of a newly founded lineage *p* may be selected for. Also, the importance of these two fitness components may change with population size, with survival probability being more important at small lineage size and net growth becoming more decisive for larger lineage sizes. We define these two measures of fitness as

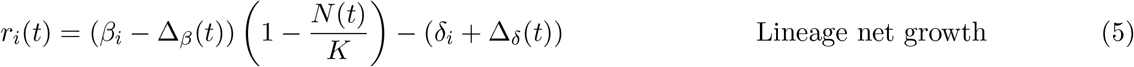

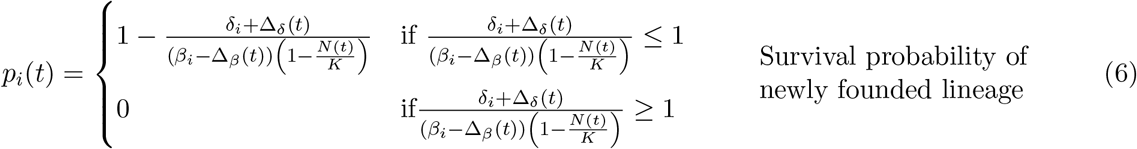

The survival probability here follows from a simplified branching process under the assumption that during the potential establishment of a mutant lineage, the population size of the remaining population will stay approximately constant (see Supplementary Section A.1). Assuming a large carrying capacity *K*, the density dependence vanishes and the survival probability becomes equal to one minus the extinction probability for newly founded lineages as derived by others (Xue and Leibler, 2017; Coates et al., 2018; Marrec and Bitbol, 2020b).

We numerically confirmed the agreement of the survival probability definition with simulations of our model for the case of no mutation (*μ* = 0) (Fig. S1). Note that the fraction of birth rate over death rate has also been proposed as a fitness measure for a model that is identical to ours, but lacks mutations (Parsons and Quince, 2007).

Adaptation will either be driven by selection for the fittest lineage in the stochastic model or determined by the fitness gradient in the deterministic model. In both cases, adaptation manifests as a changing average population trait combination. The direction of adaptation in trait space should be determined by the gradients of the two fitness components in the absence of treatment. We can compute those gradients as

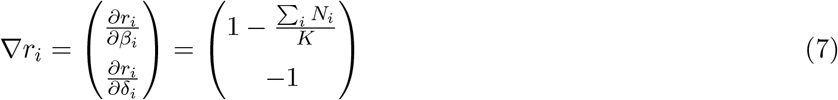

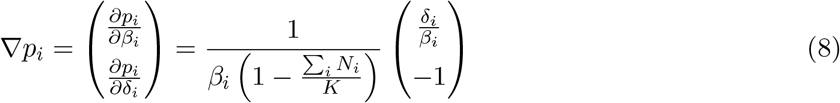

In the deterministic model (Eq. 3) we explicitly prescribe whether adaptation should follow the net growth or the survival probability fitness gradient and thus substitute *φ*(*t*) by *r*(*t*) or by *p*(*t*). If adaptation is determined by net growth we obtain

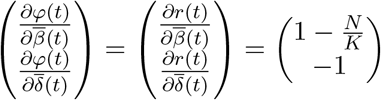

If adaptation is driven by survival probability we obtain

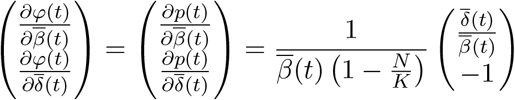

### 2.5. Treatment types

Treatment can either immediately kill part of the population or rig the chances of a population to grow by decreasing birth rates or increasing death rates (Fig. 1). The first case, which affects density directly, causes a direct, instantaneous population size reduction. The second case, which affects traits, brings about an indirect, gradual population size decline where on average more death events than birth events occur. These two treatment types thus differ in their temporal structure. Whereas the first treatment occurs instantaneously, the latter treatment is applied for a defined time span, during which the treatment alters the effective birth and death rates of cells, similar to (Marrec and Bitbol, 2020b). We assume that the density-affecting treatment type targets all cells homogeneously, irrespective of their traits. The additive trait changes during trait-affecting treatment are also equally applied to all lineages, resulting in different relative trait changes, depending on the trait values of each lineage. We represent different trait-affecting treatment types as vectors of length Δ in trait space with components given in Fig. 1. Besides the pure, static (affecting birth rates only, horizontal) or toxic (affecting death rates only, vertical) treatments, we account for the fact that the boundaries between static or toxic treatment are often blurred. The same drug can be static or toxic, depending on the dose (Masuda et al., 1977), or treatment intentionally consists of two different drug types that each act more static or toxic (Coates et al., 2018; Jaaks et al., 2022). Thus, we include a mixed treatment where both treatment vector components Δ_*β*_ and Δ_*δ*_ have the same length. Additionally, we propose two treatment types that also combine static and toxic components but additionally account for the shape of the fitness landscape. The minimizing growth treatment counters the net growth rate fitness gradient (Eq. 7) and has vector components 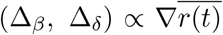 where 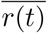 is the average net growth rate of the population at time *t*. The maximizing extinction treatment counters the survival probability fitness gradient (Eq. 8) and has vector components 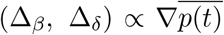 where 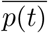 is the average survival probability of the population at time *t*. The minimizing growth treatment components are density-dependent, the maximizing extinction treatment components are trait-dependent, i.e. a function of the population average trait combination (Fig. 1).

## 3. Results

### 3.1. Trajectories of adaptation in untreated populations

When suddenly faced with challenging environments, rapidly proliferating cell populations can quickly adapt by acquiring mutations, often resulting in continuing population growth. We represent the resulting phenotypic changes as changed trait values of offspring lineages relative to the trait values of their parental lineages. Such phenotypic adaptations allow for population size increases and realize a continuously changing average population trait combination (Fig. 2). The population size increases are characterized by a succession of fitter and fitter lineages that raise the effective carrying capacity *N* ^*^ (Eq. 4), which allows the population size to increase further. Thus trait adaptation acts as a rubber band here that is extended by adaptive steps and contracts as growth closes the gap between population size and effective carrying capacity. The adaptive steps form a trait space trajectory that travels from the trait combination of the initial parental lineage to smaller death rates and larger birth rates.

**Figure 2.**
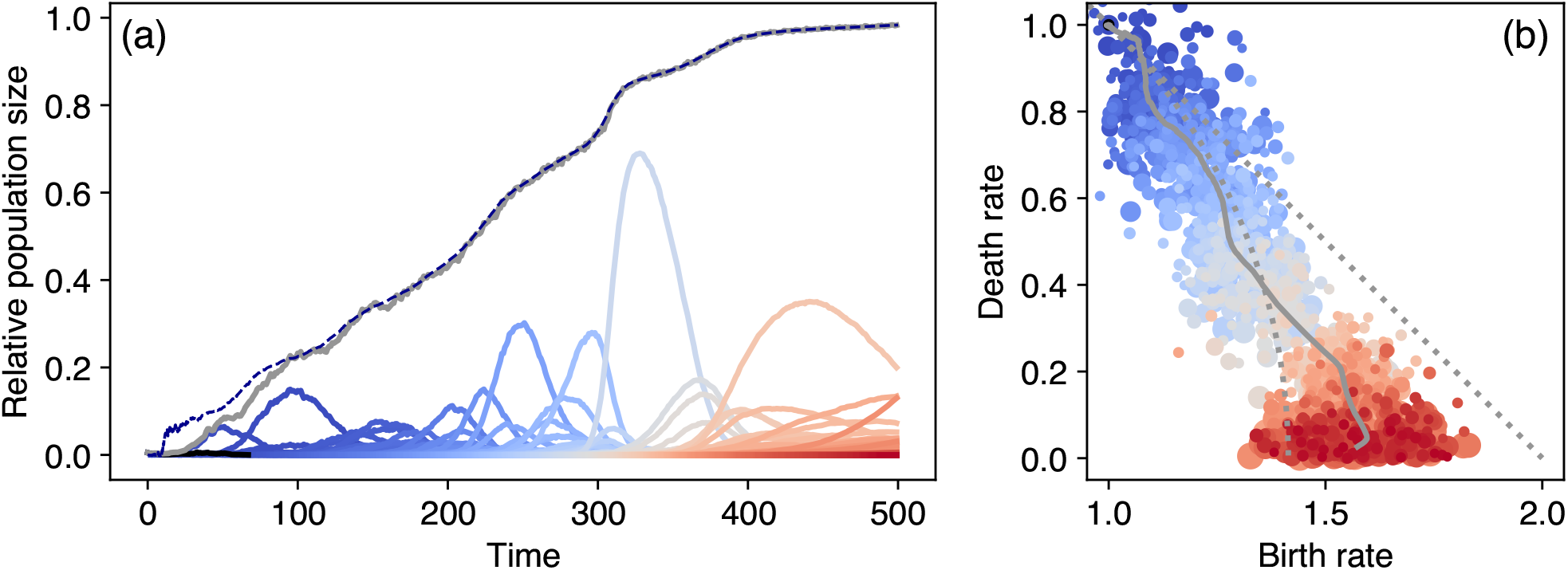
Exemplary population and trait dynamics for adaptation in challenging environments. (a) Starting from small initial numbers (*N*_0_*/K* = 0.01) the total population size (grey line) relative to the carrying capacity, *N* (*t*)*/K*, increases in a succession of fitter and fitter lineages (depicted by the blue-to-red colors indicating the order of appearance). For clarity, we show here only those lineages that persist for more than 10 time units. The dashed line shows the effective carrying capacity where population change is zero in the deterministic model (Eq. 4). The appearance of fitter lineages increases the effective carrying capacity and allows for a further increase in population size. (b) The trait combination of each lineage in panel (a) is shown here with the same color coding, with the grey line now depicting the population average. The point size is determined by the persistence time of a lineage relative to the longest persistence time. Starting from challenging conditions of birth rate *β*_0_ = 1 and death rate *δ*_0_ = 1 the population average trait combination (grey line) travels through trait space describing the trait space trajectory of adaptation. The dotted grey lines represent the net growth fitness gradient at small population sizes (straight line) and the survival probability fitness gradient (circular line).

We hypothesize that this trajectory is the outcome of the stochastic exploration of trait space that climbs up a fitness landscape, with fitter lineages out-competing less fit lineages. This fitness landscape can be characterized by fitness gradients and we propose net growth rate and survival probability as potential fitness components that generate these gradients. For our model, we see that the gradients of these two fitness components are not necessarily aligned. The vector representations of the net growth rate fitness gradient are parallel throughout trait space, indicating higher net growth rates for high birth rates and low death rates, resulting in a unidirectional, trait-independent fitness gradient (Fig. 3a). The vector representations of the survival probability fitness gradient form a circular vector field, indicating a trait-dependent fitness gradient with higher survival probability for high birth rates and low death rates (Fig. 3b).

**Figure 3.**
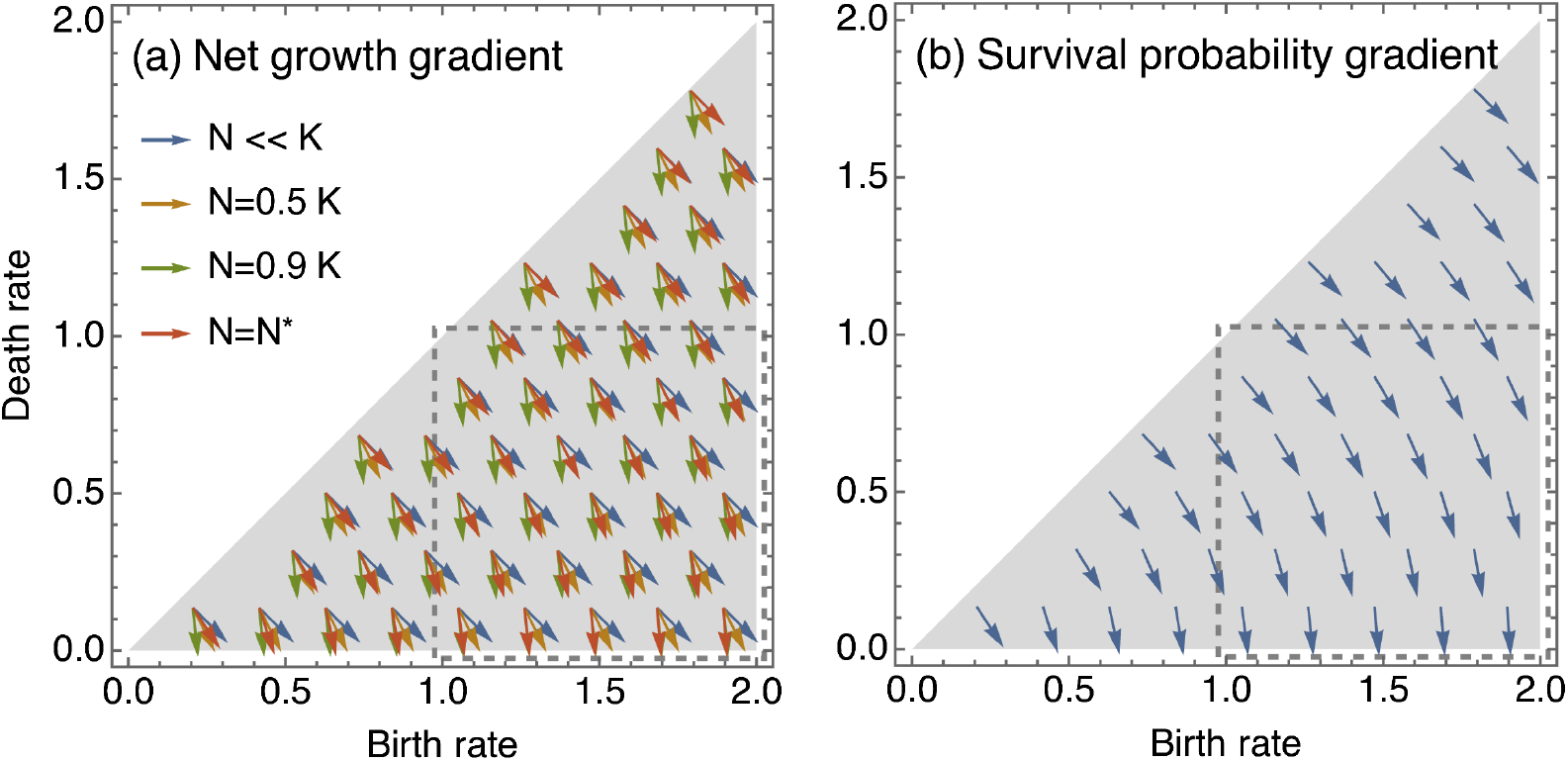
Predicted adaptation directions in trait space. (a) The direction of the net growth gradient is density-dependent, but trait-independent (Eq. (7)). (b) The direction of the survival probability gradient is density-independent, but trait-dependent and has a circular shape (Eq. (8)). At the effective carrying capacity *N* ^*^, depicted by the red arrows in panel (a), the net growth fitness gradient is parallel to the survival probability fitness gradient. Note that the effective carrying capacity depends on the traits, this causes the apparent trait dependence of the net growth gradient at effective carrying capacity. Given these gradients and initial parental lineages starting from *β*_0_ = *δ*_0_ = 1 the trait trajectories are moving mainly within the region of trait space enclosed by the grey dashed rectangle. Therefore, we zoom in on this region when visualizing trait space trajectories such as in Fig. 2.

The direction of the gradient of net growth ∇*r* is density-dependent, i.e. it changes with population size (Eq. 7). The direction of the gradient of survival probability ∇*p* does not depend on population size but is trait-dependent (Eq. 8). Interestingly, we find that both fitness gradients are parallel as soon as the manifold of zero population size change is reached and the population size equals the effective carrying capacity, *N* (*t*) = *N* ^*^ (Eq. 4, Fig. 3). Therefore, only in the initial phases of adaptation (Fig. 2a), or during and short after treatment when the population size deviates from *N* ^*^ the two fitness components may have non-parallel directions and thus differently affect the direction of adaptation steps. As soon as the total population size reaches *N* ^*^, the effects of the two fitness components cannot be disentangled, leaving us to conclude that they together dictate the trajectory of trait adaptation.

Successful adaptation in unfavourable conditions is a stochastic event. When starting with an initial wildtype population size of *N*_0_ *>* 0 and equal birth and death rate, the net growth rate is negative and the survival probability is zero (Eqs. 5, 6). Thus, the wildtype lineage inevitably goes extinct in our model, and population survival can only be achieved by adaptation and the succession of fitter lineages as described above, i.e. evolutionary rescue. The success of this adaptive process and its average trajectory can be depicted by combining a large number (1000) of independent replicates (Figs. 4, 5). We find that moving the trait combination of the first parental lineage further to the upper right corner of trait space, and thus increasing both the initial birth and death rate equally, increases the number of extinct replicate populations, indicating a lower probability of successful adaptation and evolutionary rescue. As expected, we find that a larger initial parental population and a higher mutation probability per birth event increase the rescue probability as this increases both the pool from which new lineages can emerge and rate at which they appear (Figs. 4, S2).

**Figure 4.**
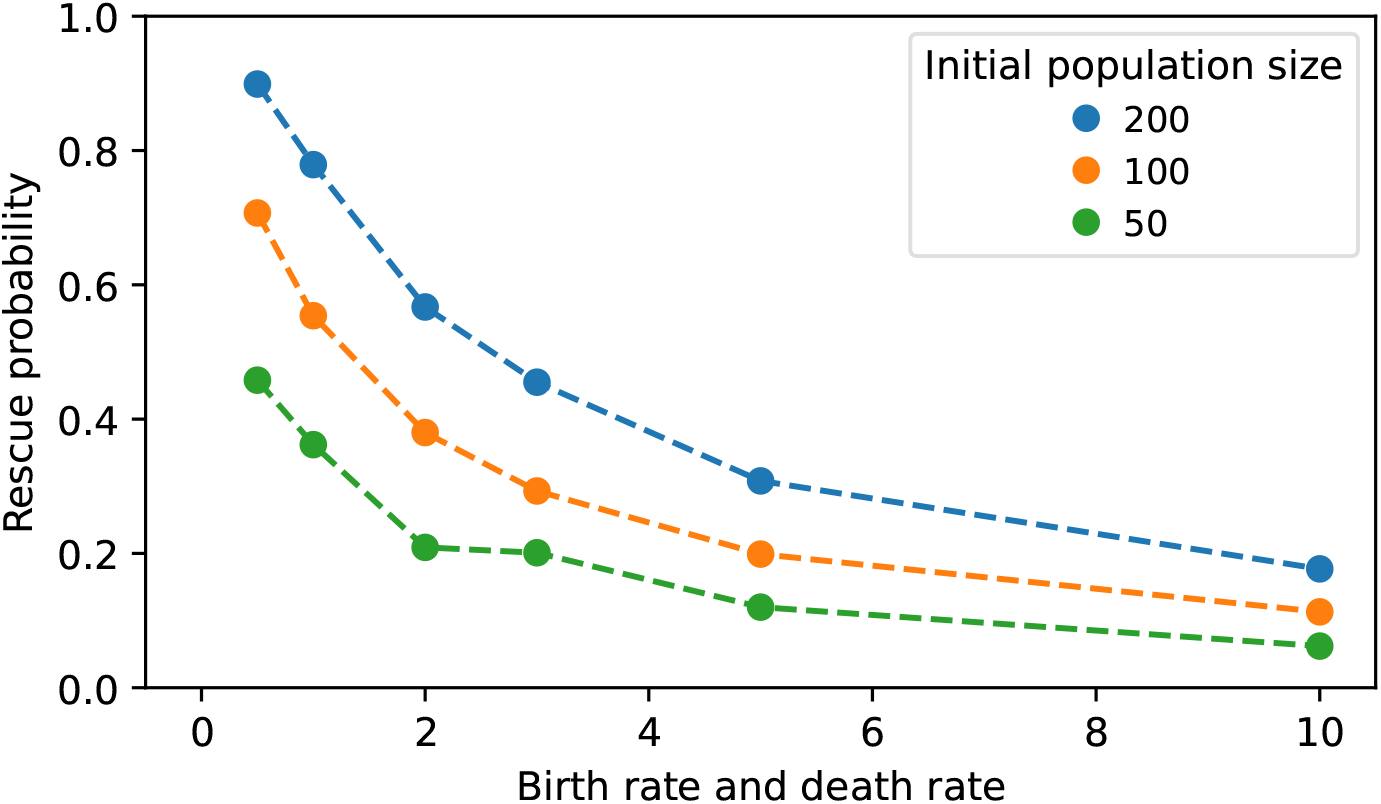
Probability of evolutionary rescue. First parental populations with higher turnover as characterized by higher levels of equal birth and death rate are less likely to successfully adapt and escape extinction. Rescue probability is here defined as the fraction of non-extinct replicate populations after *t* = 500, which allows non-extinct populations to move far into trait space regions of high net growth rate and high survival probability (see for example Fig. 2). Simulations are started from the initial parental population size *N*_0_ using 1000 replicates.

**Figure 5.**
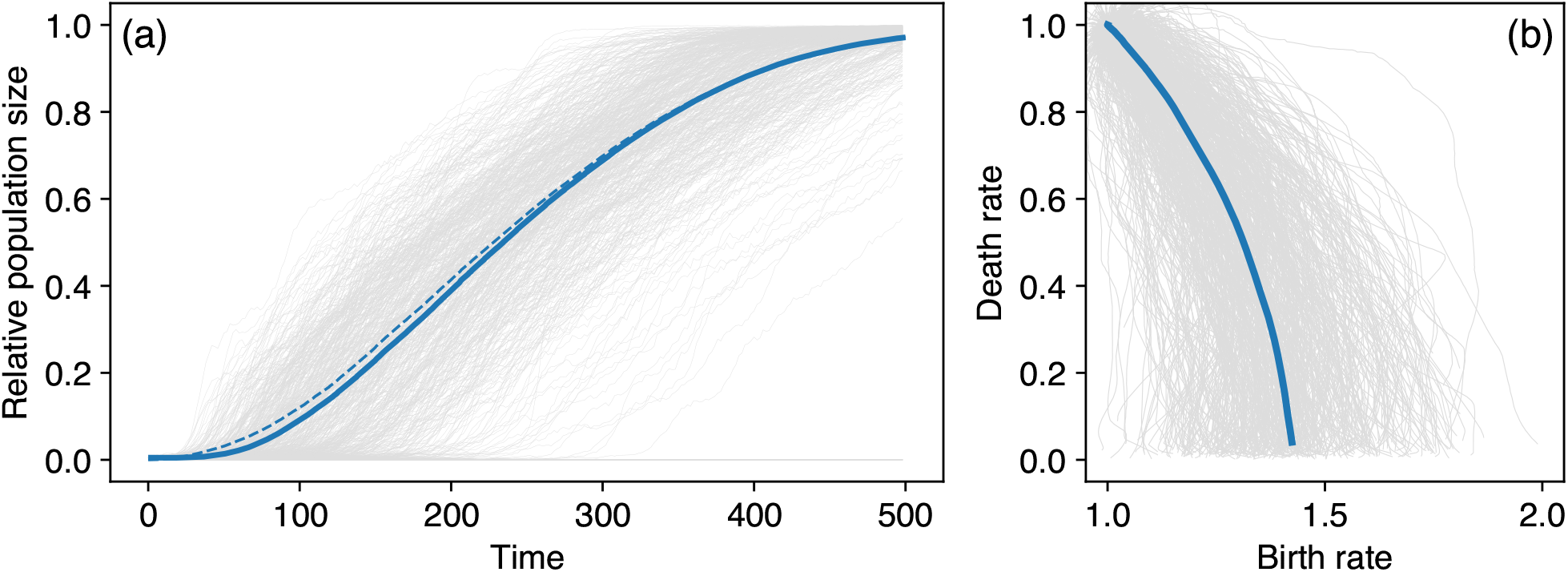
Ensemble population size dynamics and trait trajectories without treatment. (a) The population size *N* increases to the carrying capacity *K* in those replicate populations (light grey) that do not go extinct. The solid blue line represents the ensemble average of the surviving populations, the dashed blue line is the effective carrying capacity *N* ^*^ of these replicates. (b) The trait trajectories (light grey) of all replicates on average describe a circular shape (blue line). To characterize the ensemble, we consider 1000 replicates of the simulation in Fig. 2.

In those replicates where the population does not go extinct, we see that the ensemble average population size tracks the effective carrying capacity *N* ^*^ of the ensemble and approaches the carrying capacity *K* in a sigmoidal fashion (Fig. 5). The corresponding ensemble trajectory of untreated trait adaptation describes a circular shape in trait space, as predicted by both the survival probability fitness gradient and, if the population size equals the effective carrying capacity, the net growth fitness gradient.

### 3.2. Trajectories of adaptation in treated populations

For both plausible fitness gradients we can construct geometrical hypotheses about the effect of treatment on the adaptation trajectory. Visualizing the fitness isoclines (the lines of equal fitness) in trait space as the rectangular bases for the fitness gradient vectors helps to work out this effect (Fig. 6). We consider treatment types that either target the population size directly, or indirectly by additively shifting the traits of the cells which subsequently decreases population size. Both the direct as well as the indirect effect on population size induce a density-mediated rotation in the net growth fitness isoclines (Fig. 6a). This causes a less vertical predicted adaptation direction with a larger birth rate component from the net growth fitness component. The trait-affecting treatment types temporally displace the population in trait space but have no direct effect on the net growth fitness component due to the parallel fitness isoclines (Fig. 6b). Similarly, the survival probability fitness component is independent of population size and thus not affected by density changes (Fig. 6c). However, when the population is displaced in trait space the circular shape of the survival probability fitness component changes the predicted adaptation direction to become less vertical under trait-affecting treatment (Fig. 6d). Thus, we hypothesize that both treatment types would drive less vertical adaptation trajectories.

**Figure 6.**
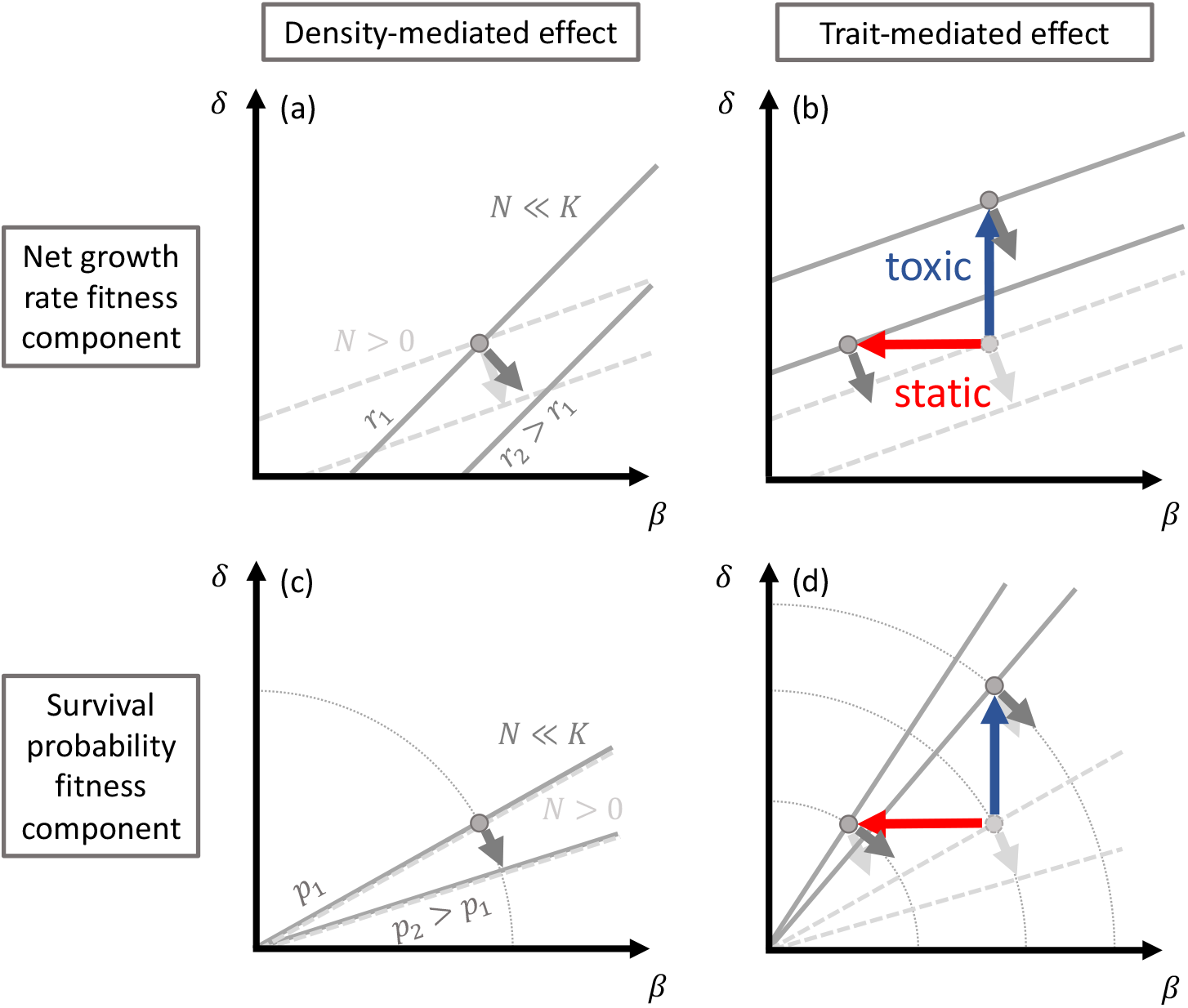
Density-mediated and trait-mediated treatment effects predict less vertical trait adaptation trajectories. The fitness isoclines (contours of equal fitness in trait space) are by definition perpendicular to the fitness gradient vectors for a given point in trait space. Fitness isoclines in the absence of treatment are depicted by dashed grey lines, fitness isoclines affected by treatment are shown as solid, dark grey lines. Similarly, realized trait combinations that include the effect of treatment are shown by dark grey points. They deviate from their light grey, untreated counterparts in the case of trait-affecting treatment. Potential changes in the adaptation direction are indicated by a difference between untreated (light grey) and treated (dark grey) fitness gradient vectors, and corresponding fitness isoclines with different angles relative to the axes.

We investigate the effect of treatment on the adaptation trajectory by periodically applying the different treatment types on populations that grow from small population sizes and ascend the fitness gradient (Fig. 7). If the replicate populations escape extinction, they increase in population size and reach the carrying capacity *K*. The density-affecting treatment type reduces the population size of each lineage by a bottleneck factor *f*. This decreases competition and allows surviving lineages to achieve higher net growth rate. This competitive release causes the population size to recover to higher levels after the first treatments than in the untreated control (Fig. 7a). However, newly established, fitter lineages are especially prone to extinction when the bottleneck treatment reduces lineage sizes to small fractions, which limits the exploration of trait space and hinders a rapid adaptation towards faster net growth rates and higher survival probabilities. Therefore, the populations that undergo stronger bottleneck treatments approach the carrying capacity slower and have shorter trait trajectories (Fig. 7a,b). The trait-affecting treatment types also show the competitive release pattern of recovery to population sizes higher than the untreated control. Here, the population sizes repeatedly recover to higher values after treatment and the carrying capacity is approached faster than in the untreated control (Fig. 7c,d). Similar to the untreated population size time series, also under treatment the population size is tracking the effective carrying capacity *N* ^*^. We find that the trait trajectories of treated populations deviate from the untreated controls as predicted from our geometrical hypotheses (Fig. 6). We observe that the deviations are caused by more horizontal adaptation steps right after the density-affecting treatment or during the trait-affecting treatment (Figs. S7, S8). This results in longer adaptation trajectories that are elongated towards higher birth rates. The traits change in a step-wise pattern over time for the density-affecting treatment, with large adaptive steps immediately after the treatment time points (Fig. 8a). Trait-affecting treatment increases the rate of trait change which results in a ramp-like pattern of the traits over time Fig. 8b).

**Figure 7.**
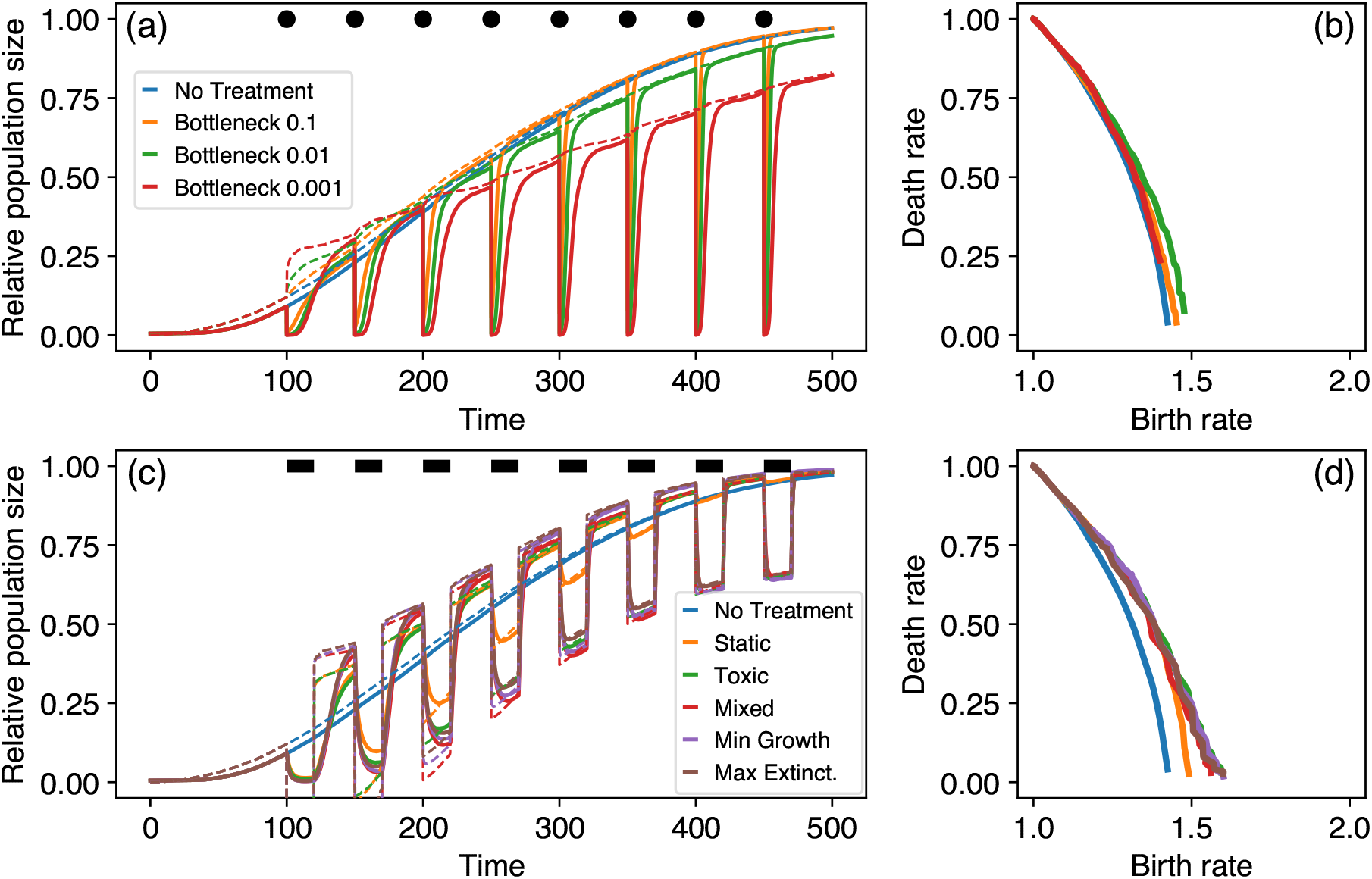
Ensemble population size dynamics and trait trajectories under treatment. (a) Density-affecting treatment applies regular bottlenecks and instantaneously decreases the population size of each lineage to a small fraction at time points indicated by the black points. The treatment strength is varied by decreasing the remaining fraction of each lineage after treatment (different colors). The population size dynamics track the effective carrying capacity (Eq. 4, dashed lines). (b) The density-affecting treatment affects the ensemble trait trajectory by triggering sudden trait changes. (c) Trait-affecting treatment types result in prolonged phases of reduced population size (indicated by the black bars). Again, the dashed lines depict the effective carrying capacity dynamics. (d) The ensemble average trait trajectories under trait-affecting treatment deviate from the no treatment reference and reach higher birth rates. Exemplary population size time series and trait trajectories for bottleneck, static and toxic treatment are shown in Figs. S4-S6. As before we performed 1000 replicate simulations and computed ensemble averages from the surviving replicates.

**Figure 8.**
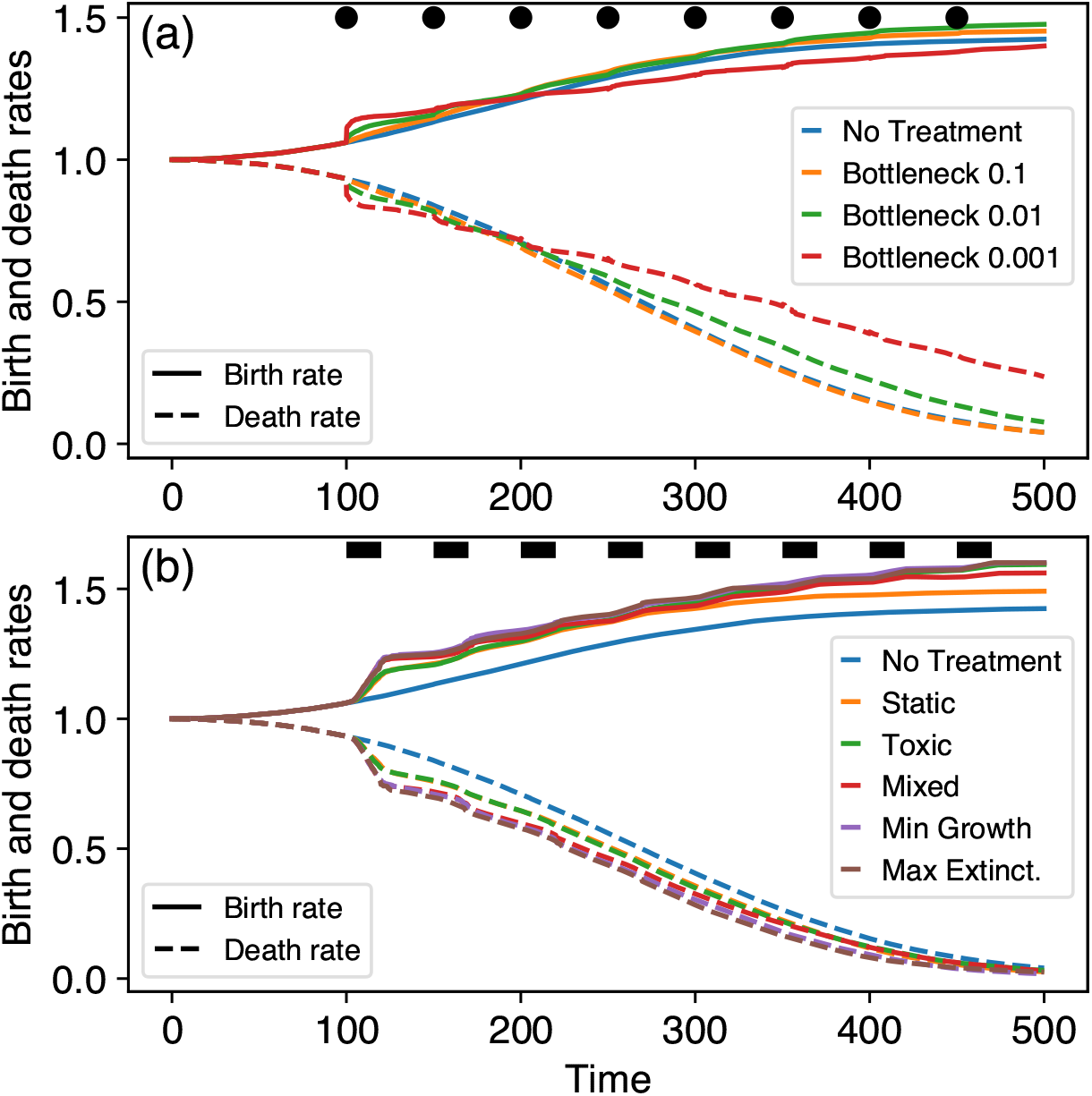
Trait dynamics under treatment depict the speed of adaptation. (a) Density-affecting treatment causes short spikes in adaptation speed that manifest in step-wise changes of the ensemble trait average. (b) Trait-affecting treatment temporarily accelerates the changes in ensemble trait averages leading to ramp-like trait changes. The different colors refer to the treatment types, the solid and dashed lines represent birth and death rates, respectively.

We find that the dynamics of those trait-affecting treatment types that contain toxic components are similar both in the population size and the trait dynamics. The purely static treatment, however, differs considerably. As the population size approaches the carrying capacity, the effect of the static treatment is reduced as its net growth reduction is density dependent and proportional to 1 − *N/K* (Eq. 1). This manifests in decreasing density reductions during treatment phases (Fig. 7c). Accordingly, after similar initial trajectories, the adaptation trajectory under purely static treatment later deviates from the adaptation trajectories for the other treatment types that contain also density-independent toxic components (Fig. 7d). We observe similar patterns also in the deterministic description of the adaptive process using a quantitative genetics approach where we explicitly specify the gradient of trait adaptation (Eq. 3, Figs. S9, S10).

These fundamental effects of different treatment types on population dynamics and trait adaptation trajectory translate to treatment efficiency and the possibility for the populations to escape the treatment, i.e. evolve treatment tolerance. In our model and for the chosen parameters, approximately half of the replicates go extinct without any treatment due to stochastic extinction in the initial phases of adaptation. This pattern is caused by the initially equal birth and death rates. Equal birth and death rates imply zero net growth and thus inevitable extinction due to stochastic population size fluctuations. The adapting populations depart from this. Applying treatment increases the fraction of extinct replicates, which we use as a measure to quantify the treatment success rate (Fig. 9). As expected, a higher treatment strength that removes a larger proportion of cells per lineage increases the success rate of the density-affecting treatment type. Among the trait-affecting treatment types, pure static and toxic treatments achieve a similar success rate. Interestingly, combining static and toxic treatment components results in a considerably higher success rate. Here, the success rate of treatment types that counter either the net growth fitness gradient or the survival probability fitness gradient is slightly higher than the ‘Mixed’ treatment type that non-adaptively blends the static and toxic components in equal proportion.

**Figure 9.**
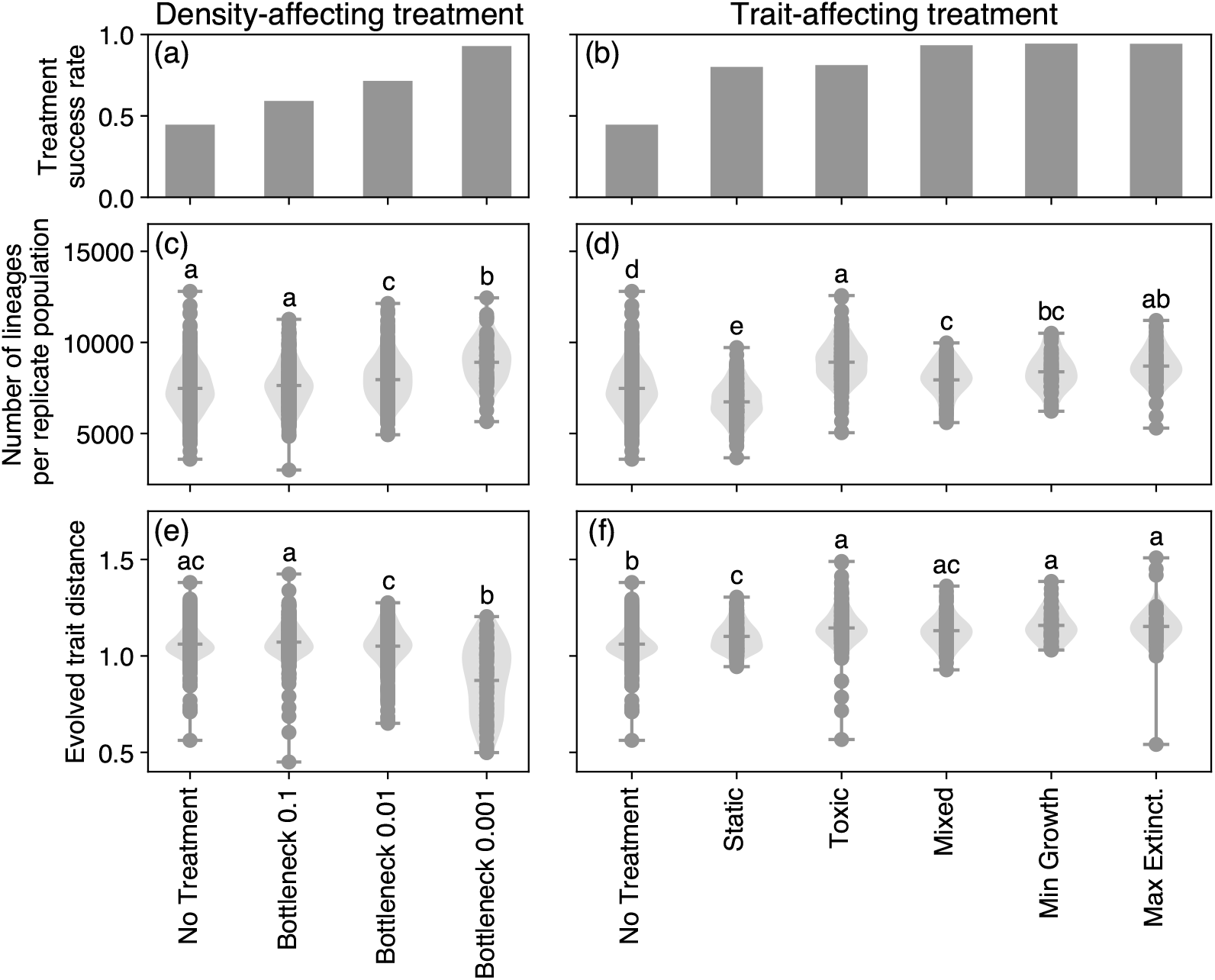
Treatment effects for density-affecting (left column) and trait-affecting (right column) types. Panels (a) and (b) show treatment success rate measured as the fraction of extinction among the 1000 replicate populations at *t* = *t*_*f*_. Panels (c) and (d) show the number of lineages that have been created by mutations in each non-extinct replicate population. Panels (e) and (f) show the distance between the first parental trait combination and the last average trait combination of each non-extinct replicate population. In panels (c)-(f), the same lower-case letters above two treatments indicate that the two sets of data points could have been generated from the same underlying distribution. Differing lower-case letters thus indicate differences between treatments. Unique letters indicate treatments that are statistically different from all other treatments. The grouping into statistically different groups was determined using the Tukey’s HSD implementation from the statsmodels module (v0.13.0) in Python 3.8 and assigned with the pairwisecomp letters function written by Philip Kirk (https://github.com/PhilPlantMan/Python-pairwise-comparison-letter-generator). A treatment can be part of multiple groups by being indifferent to each one of them and thus receive multiple letters.

An interesting pattern emerges for the overall number of lineages that are eventually created during the adaptation from one parental lineage, which relates to the evolutionary potential of the population. We find that treatments that particularly increase mortality while not decreasing birth rates lead to a higher number of created lineages. The higher mortality decreases the density limitation of birth rates, which enables high net birth rates and accordingly high mutation rates. Particularly the stronger density-affecting treatments and the purely toxic treatment result in the creation of more mutant lineages. Whether these lineages are expanding successfully and thus shift the population average trait combination depends on the survival of the newly created lineages. Accordingly, we find a reduced exploration for the strongest density-affecting treatment measured by the distance between the first parental trait combination and the population average trait combination at the end of our simulations. For the trait-affecting treatment types, we find an opposite correlation. Here, more lineages also enable a further trait space exploration. Newly created lineages are in general more endangered by extinction than established lineages, simply because of their smaller cell numbers, which makes a stochastic crossing of the extinction boundary more likely. During bottleneck treatment the relative effects of treatment on the extinction risk for newly created, fitter lineages versus established, less fit lineages are equal, whereas the absolute effects are different as it is more likely for small lineages to be driven to population sizes below a single cell. During trait-affecting treatment, the relative effect of treatment is smaller for smaller, but fitter lineages than for established, less fit lineages, whereas the absolute effects are equal. This may explain the observed differences in the correlation of number of lineages and evolved trait distance. It is interesting to note that treatments with higher success rate were also found to induce faster trait changes (Fig. 8), pointing out a potential trade-off of treatment success versus driving tolerance evolution.

### 3.3. Which fitness component is more important?

We found that treatment types that counter the potential fitness gradients achieve the highest success rates. However, we have not conclusively answered whether the net growth fitness gradient or the survival probability fitness gradient are more decisive for the eco-evolutionary dynamics in our model. To gather more evidence on this, we sampled the initial adaptation direction from different initial trait combinations to visualize the realized fitness gradient that acts on the adapting populations in trait space (Fig. S11). We indeed find that the realized fitness gradients are non-parallel in trait space, indicating that for larger birth rates and smaller death rates adaptation is driven by decreasing death rate, and increasing birth rate becomes less important. The visual similarity of this pattern to the survival probability fitness gradient hints at a larger importance of the survival probability fitness gradient at first glance. However, also the net growth rate becomes larger for larger birth rates and smaller death rates, which speeds up the population size increase during the short observation window of initial adaptation. Because of the density-dependence, these larger population sizes turn the net growth fitness gradient to be more vertical (see Fig. 3a). Also, we observe that the initial adaptation direction is largely parallel along the diagonals in trait space, which correspond to the net growth fitness isoclines for small population sizes, which favours the net growth fitness gradient to be more important.

To investigate whether the differences in initial adaptation direction are indeed caused by the densitydependence of the net growth fitness gradient, we again investigated the initial steps of adaptation with parameters that minimize the density change within our observation window. We decreased the initial population size and time span and increased the carrying capacity and find that the adaptation direction indeed becomes more horizontal, indicating a larger importance of the net growth fitness gradient than the survival probability fitness gradient. If the survival probability fitness gradient would be predominantly driving the adaptation, we would expect that the initial steps of adaptation change along the net growth fitness isoclines (except for the diagonal passing through the origin) and we would not expect a density dependence.

In the deterministic model (Eq. 3), we are explicitly prescribing the fitness measure that determines the direction of trait adaptation. If we choose the net growth as the determining fitness measure we find trait trajectories that change with treatment and reproduce the trajectories obtained from simulations (Fig. S9). However, if we set the survival probability as the determining fitness measure in the deterministic model the trait trajectories under density-affecting treatment do not deviate from the trajectories without treatment, thus contrasting the observation in the simulations (Fig. S10). Therefore, more evidence points towards net growth rate maximization as the determinant of trait space adaptation trajectories in our simulations, even though we cannot falsify that the survival probability fitness gradient could also play an important part.

## 4. Discussion

During the onset of cancer establishment and the spread of pathogens from a chronic infection, populations of small size have to break with homoeostatic regulations that aim to prevent their expansion. Adaptation by trait evolution allows them to climb up the fitness landscape and eventually escape stochastic extinction, that would be unavoidable without adaptation. In this study, we reduced the complexity of cancer cells and pathogenic bacteria to the three basic processes of birth, death and mutation, and investigated i) the shape of the fitness landscape, ii) the adaptive trajectories of trait evolution and iii) how these trajectories are altered by treatment. We proposed net growth rate and survival probability as possible fitness measures that are increased by evolution. We found that both of these measures motivate a circular adaptive trajectory in the trait space spanned by birth rate and death rate (Fig. 3). Indeed, this circular trajectory is recovered in stochastic simulations (Fig. 5) and altered by treatment in agreement with geometrically derived hypotheses (Figs. 6, 7). Interestingly, we find that adaptive steps that maximize net growth rate or survival probability always have parallel components, indicating no strong conflict between optimizing for either of the two plausible fitness measures.

In this study, we deliberately chose parameters that would result in occasional extinction of replicate populations to represent the stochastic nature of the establishment of cancer or bacterial infections and the stochasticity in treatment response (Coates et al., 2018; Alexander and MacLean, 2020). This results in a setting where evolutionary rescue is required for the populations to prevent their extinction. In our model, the population dynamics are captured by the dynamics of the effective carrying capacity which is the target population size that the total population size is tracking over time. If birth rates and death rates are equal, the effective carrying capacity is zero and the population goes extinct deterministically. The effective carrying capacity becomes positive only if the death rate becomes smaller than the birth rate by trait adaptation, thus also increasing the chances of population establishment.

The shape of the fitness landscape has important implications for the effect of turnover on the rescue probability in the cancer or bacterial cell population, which we can again address using geometrical arguments of the fitness isoclines. A faster turnover implies that birth rates and death rates of the associated cells are higher. This puts fast-turnover cells in the upper right corner of our birth-death trait space, and slow-turnover, quasi-dormant cells in the lower left. The circular fitness gradient vector field of survival probability implies radial fitness isoclines, resulting in an increasing distance between isoclines going from slow to fast turnover. Therefore, the same adaptation step in trait space gains a smaller increase in the survival probability of fast-turnover cells than in slow-turnover cells. This implies that evolutionary rescue is less likely in populations of fast-turnover cells, which we indeed find when comparing the fractions of surviving replicates for different initial parental trait combinations of equal birth and death rates. Interestingly, Kuosmanen et al. (2022) come to similar conclusions in a slightly different model. Importantly, this pattern can be affected by the assumptions on the mutational effect sizes. Throughout this study we have assumed additive mutational effects where the adaptation step sizes are independent of the trait values. If, however, the mutational effects were dependent on the trait values, as for example in the case of multiplicative mutational effects (Fig. S3), this pattern will change. Accordingly, we find that multiplicative mutational effects compensate for the increasing distance of radial fitness isoclines at larger birth and death rates and the rescue probability becomes largely independent of turnover.

Besides the shape of the fitness landscape, the declining rescue probability for faster turnover may also be explained with the higher rate at which the initial parental lineage declines. At equal birth and death rate, the logistic competition term results in a deterministic rate of population decline of −*β*_0_ *N*_0_(*t*)^2^*/K* in our model, which increases proportional to the birth rate. As this initial parental lineage is the source from which offspring lineages are created, a faster decline shortens the time window during which fitter lineages can emerge and impedes the race against extinction (Orr and Unckless, 2008, 2014; Carlson et al., 2014; Marrec and Bitbol, 2020a). On the other hand, in fast-turnover cells mutations occur more frequently because of the higher birth rate, which could speed up the ascend of the survival probability fitness gradient. Our results show that the higher realized mutation rate cannot compensate for the two detrimental effects of faster turnover firstly requiring larger trait changes for the same gain in survival probability and secondly leading to a faster decline in the initial parental lineage.

Cancer cell populations as well as bacterial biofilms in chronic infections possess a considerable genotypic and phenotypic heterogeneity (Caiado et al., 2016; Gay et al., 2016; Winstanley et al., 2016; Dhar et al., 2016). In a heterogeneous population consisting of lineages with different turnover but individually equal birth and death rates our results imply that those lineages with smaller turnover would persist longer. Evolutionary rescue would thus be achieved on average from those lower-turnover lineages hinting at a selective advantage of low turnover in heterogeneous populations in challenging environments, which may explain the therapeutic challenges posed by dormant subpopulations both in cancer (Yeh and Ramaswamy, 2015; Ammerpohl et al., 2017) and bacterial infections (Wood et al., 2013). Birth (proliferation) and death (apoptosis) are partly interlinked in their regulation (Alenzi, 2004) and measuring their rates in eukaryotic cells is possible in vitro and in vivo (Lyons and Parish, 1994). Different tissue types were shown to have intrinsically different turnover rates (Sender and Milo, 2021) and turnover can be altered experimentally (Casey et al., 2007). Several studies reported a positive correlation of proliferation and apoptosis in breast cancer (de Jong et al., 2000; Liu et al., 2001; Archer et al., 2003), which suggests a positive correlation of birth and death rate. Prognosis was found to be worse for higher birth rate (Liu et al., 2001). Our model proposes that such aggressive, quickly growing tumours with a high cell death rate are actually less likely to persist than tumours with lower turnover as the probability for evolutionary rescue decreases with turnover. This apparent dichotomy indicates that the evolutionary rescue probability of a tumour not necessarily translates into its prognosis and that clinically we tend to only observe the few high-turnover tumours that have managed to escape homeostatic regulation, while remaining blind to those with lower turnover. Also in the context of chronic bacterial infections there exist methods to assess turnover in bacterial pathogen populations in vitro (Stewart et al., 2005; Wang et al., 2010). They are currently developed for in vivo settings (Myhrvold et al., 2015; Mahmutovic et al., 2021) and will soon elucidate the different intrinsic birth and death rates of bacterial strains and species, sometimes even working out spatial parameter heterogeneity within the body (Gillman et al., 2021). It will be interesting to see whether indeed lower-turnover regions of the birth-death trait-space are found to be more populated and whether trait evolution indeed proceeds along the circular trajectory predicted by our model.

Fitness landscapes of mutational changes can be constructed from data (Watson et al., 2020) and used in treatment via evolutionary steering (Nichol et al., 2015; Acar et al., 2020). Accounting for their temporal variability (e.g. under the effect of treatment), then sometimes referring to them as fitness seascapes, has important consequences for the understanding of adaptation, such as resistance evolution (Lässig et al., 2017; King et al., 2022). For example, Hemez et al. (2020) found in a simulation study that the drug mode of action (bacteriostatic vs. bactericidal) was changing the shape of the fitness landscape. In line with this, we have found that both density-affecting and trait-affecting treatment types alter trait adaptation trajectories. The density-mediated effect of treatment rotates the fitness landscape, the trait-mediated treatment effect relocates populations to other trait combinations in trait space. Both of these effects increase the birth rate component of adaptive steps which causes treated trait adaptation trajectories to depart from untreated trajectories.

We found profound patterns of competitive release in the population dynamics of successfully adapting populations (Wargo et al., 2007). In the off-treatment phases, the treated and non-extinct populations quickly recover to population sizes up to twice as large as in the untreated reference. The competitive release is particularly strong for the trait-affecting treatment types. This is in line with the fact that the trait-affecting treatment exerts a higher relative penalty on less fit lineages than on fitter lineages as we assumed additive treatment effects and thus the mortality during treatment is higher for less fit lineages. In our model the effect of static drugs decreases as the population size approaches the carrying capacity where the effective birth rate tends to zero even without treatment and thus can not be reduced further by treatment. Contrastingly, the sustained mortality exerted by toxic treatment also at population sizes close to the carrying capacity leads to a continuing competitive release. This creates additional transient phases of population recovery after treatment phases during which birth and mutation rates are high, resulting in faster adaptation. This potentially negative effect of toxic treatment is in agreement with findings by Anttila et al. (2019) and Marrec and Bitbol (2020b) and similar to the paradoxic negative effects of apoptosis during tumour development (Labi and Erlacher, 2015). This finding also resonates with the rational behind tumour containment treatment strategies that aim at preserving sensitive subpopulations as competitors, and thus suppressors, of resistant subpopulations (Gatenby et al., 2009; Viossat and Noble, 2021).

Time-resolved surveillance of treatment responses in both cancer and bacterial infections promises to prevent resistance evolution, but is technically and practically challenging. Accordingly, the quest for personalized, resistance-proof treatment approaches remains one to be fulfilled. In a recent paper, we found that increasing the temporal frequency of surveillance has diminishing returns and also more coarse-grained surveillance patterns could achieve large treatment improvements (Raatz et al., 2021). Interestingly, in the present study we find that the mixed treatment which is agnostic to real-time information performs almost as good as the treatment types that counter the fitness gradient and thus necessitate ongoing temporal information on the population trait average. This again suggests that large treatment improvements can be achieved already with low surveillance effort. The high efficiency of static and toxic treatment combinations is in agreement with theoretical predictions (Lorz et al., 2013) and recently explored approaches in cancer treatment, such as the combination of navitoclax, a drug that increases the apoptosis rate, and cytostatics such as gemcitabine or brentuximab which decrease the birth rate (Cleary et al., 2014; Ju et al., 2016; Montero and Letai, 2018). Also in bacteria, recent findings suggest that a combination of bacteriostatic drugs (or nutrient deprivation) and bactericidal drugs indeed increase the extinction probability of bacterial microcolonies (Coates et al., 2018). However, awareness of the mechanisms of action and the interactive effects is essential, as treatment efficiency can also be reduced in combination treatments, for example if the bactericidal drug relies on cell growth that is reduced by the bacteriostatic drug (Bollenbach et al., 2009; Bollenbach, 2015; Coates et al., 2018). An additional advantage of combination therapies that was not considered in our study is that resistance is less likely to evolve in parallel against two independently active drugs. Consequently, drug interactions have important consequences not only for treatment efficiency but also for resistance evolution (Roemhild et al., 2018; Roemhild and Schulenburg, 2019; Batra et al., 2021; Jaaks et al., 2022).

In this study, we have abstracted from the physiological details of different adaptation pathways in evolving cell populations and the molecular mechanisms of the drugs used to counter these adaptations. By mapping these details to traits with clear eco-evolutionary consequences we achieved an understanding of the adaptation dynamics, identified relevant fitness components and showed the high efficiency of trait-aware treatment strategies. Current experimental and diagnostic advancements enable the identification of traits, such as birth and death rates, at realistic scales to allow for a translation between mechanistic models such as ours and experimental and clinical observations. This will further the understanding of the eco-evolutionary mechanisms at play in the dynamics of cancer and bacterial infections and sprout improved, personalized and adaptive treatment strategies.

## Data, script and code availability

The code to reproduce all figures has been deposited at https://doi.org/10.5281/zenodo.6656842. The simulation data is available at https://doi.org/10.5281/zenodo.6656847.

## Acknowledgements

We thank Hildegard Uecker for discussions and advice on the probability of evolutionary rescue. We are grateful for biological insights into the birth and death of bacteria from Javier Lopez Garrido and Alan Derman and for discussions related to birth and death of cancer cells with Susanne Sebens and Lisa-Marie Philipp. This preprint has been peer-reviewed and recommended by PCIEvolBiol (https://doi.org/10.24072/pci.evolbiol.100555).

## A. Supplement

### A.1. Derivation of survival probability fitness component

Recently, Xue and Leibler (2017) derived the extinction risk for a population founded by a small number of individuals. Their model contained also a density-dependent death rate, which makes it slightly different from ours. They set up a master equation and solved it with a generating function approach. For a single initial individual with birth rate *β* and death rate *δ* they obtain a density-independent extinction risk of

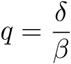

from which the survival probability for a new lineage follows as

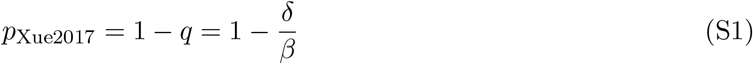

Assuming that changes in the population size of the parental lineage are small on the time scale during which the fate of a mutant is decided, i.e. whether it escapes extinction from stochastic drift or not, allows us to fix the total population size to its value when the mutant occurred at time *T*. Thus, we can include the density dependence of our model in the survival probabilty (Eq. S1) by substituting 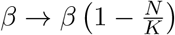. This results in a density-dependent survival probability

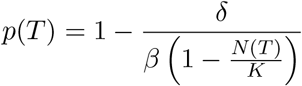

Including trait-affecting treatment effects and restricting the survival probability to the range between zero and one results in Eq. 6.

A similar derivation uses branching process techniques and arrives at an integral for the fixation probability of a mutant individual on the background of the parental population (Uecker and Hermisson, 2011)

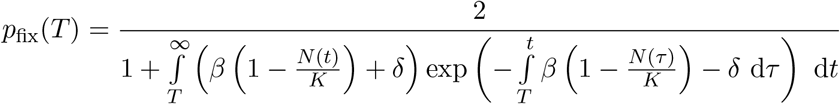

Using the same assumption of *N* (*t*) = *N* (*T*) = const. as above, this reduces to

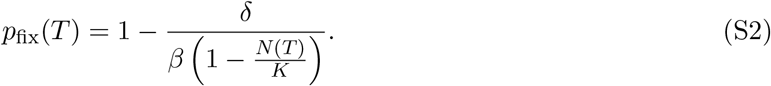

## A.2. Supplementary Figures

**Figure S1.**
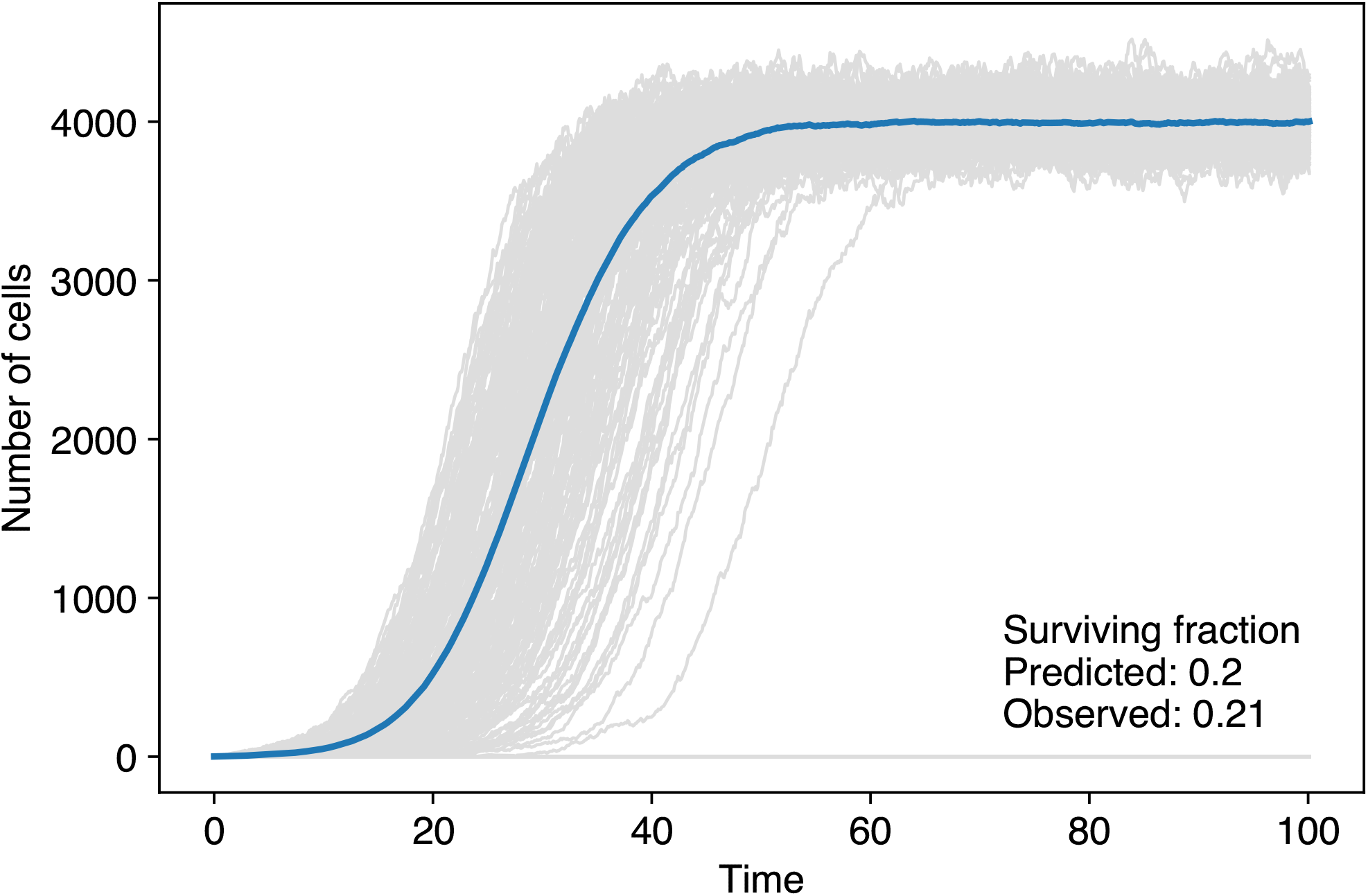
Numerical simulations of a birth-death process without mutation (*μ* = 0). Starting from *β*_0_ = 1.25 per time unit and *δ*_0_ = 1.0 per time unit we find good agreement of the observed survival probability with our survival probability definition. Grey lines are individual replicates, the blue line is the average over the surviving replicates. We used 1000 replicates, *dt* = 0.1, *N*_0_ = 1 and *K* = 20000.

**Figure S2.**
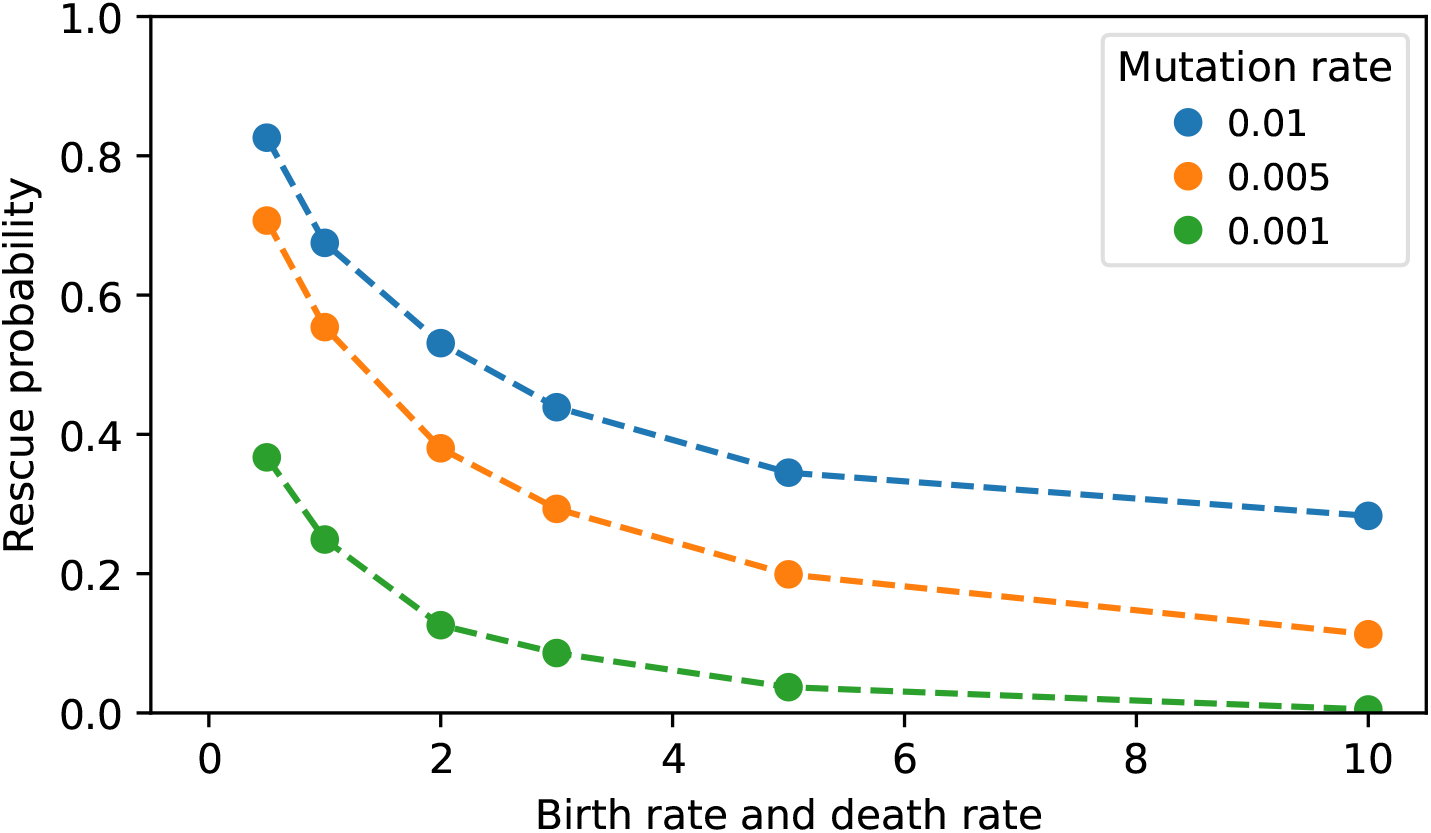
Effect of mutation rate on the probability of evolutionary rescue. Smaller mutation rates reduce the rescue probability. Plot parameters are identical to Fig. 4.

**Figure S3.**
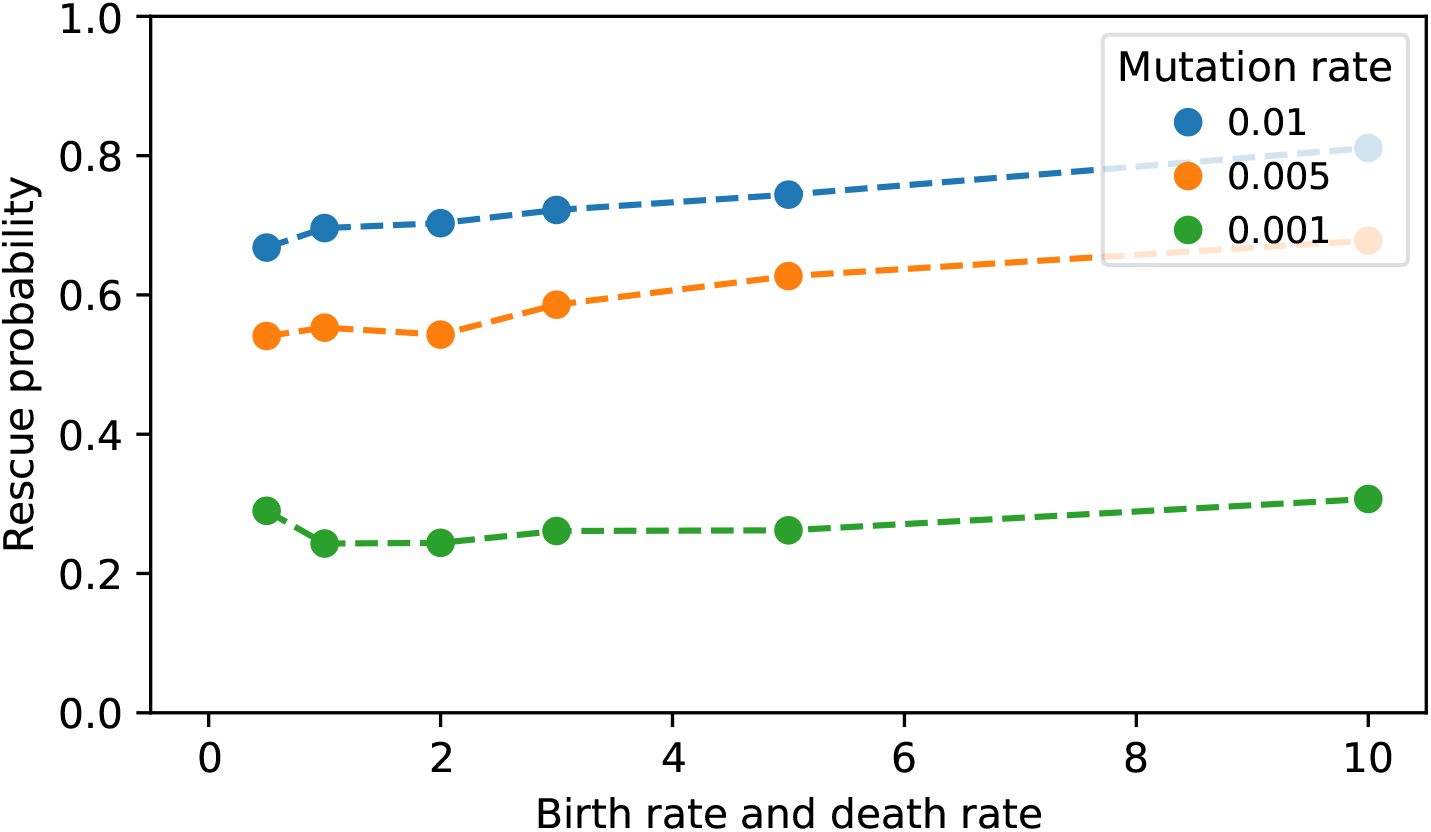
Probability of evolutionary rescue (multiplicative mutational effect). Parallel to Fig. S2 we tested the effect of multiplicative mutational effects on birth an death rates. The mutant lineages’ birth rates here are determined by *β*_mutant_ = *β*_parental_ (1 + *s*), *s* ∼ *N* (0, *σ*), and death rates are independently determined as *δ*_mutant_ = *δ*_parental_ (1 + *s*), *s* ∼ *N* (0, *σ*). Under these assumptions, the rescue probability of initial parental populations is largely independent of turnover.

**Figure S4.**
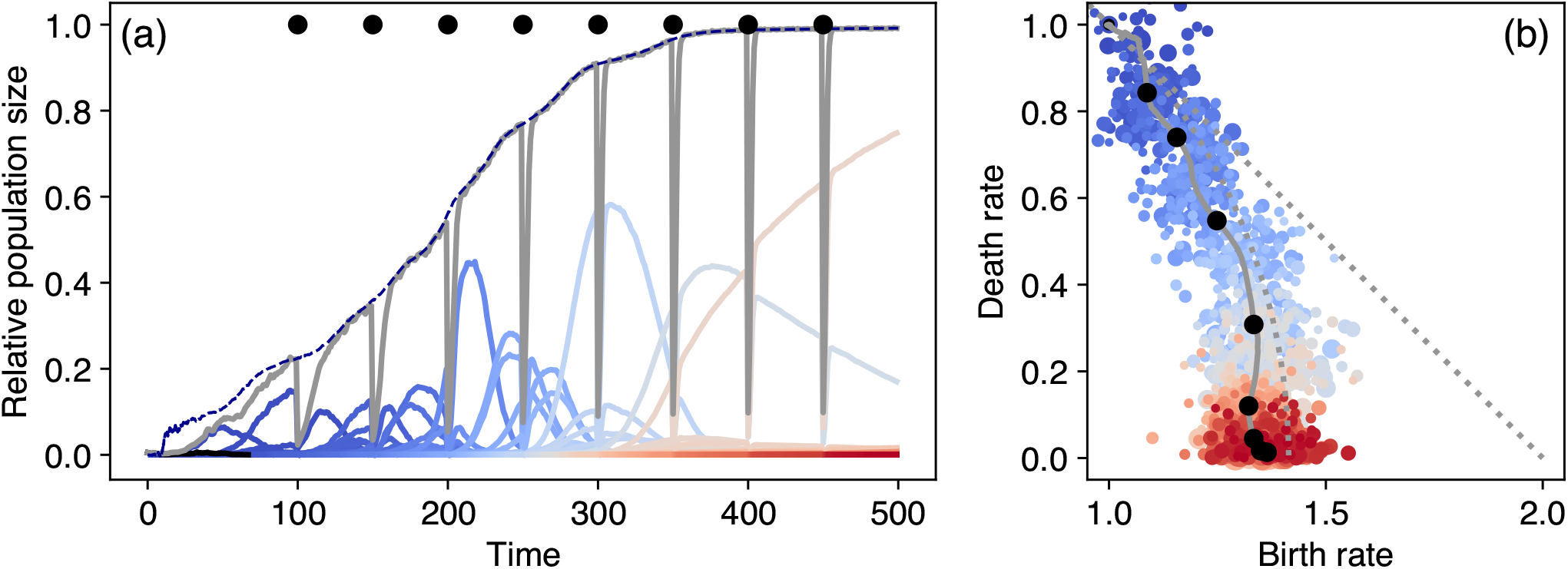
Exemplary dynamics for bottleneck treatment. Plot details and parameters as in Fig. 2. Black dots depict the times when the bottleneck instantaneously reduces the population size by a factor *f* = 0.1.

**Figure S5.**
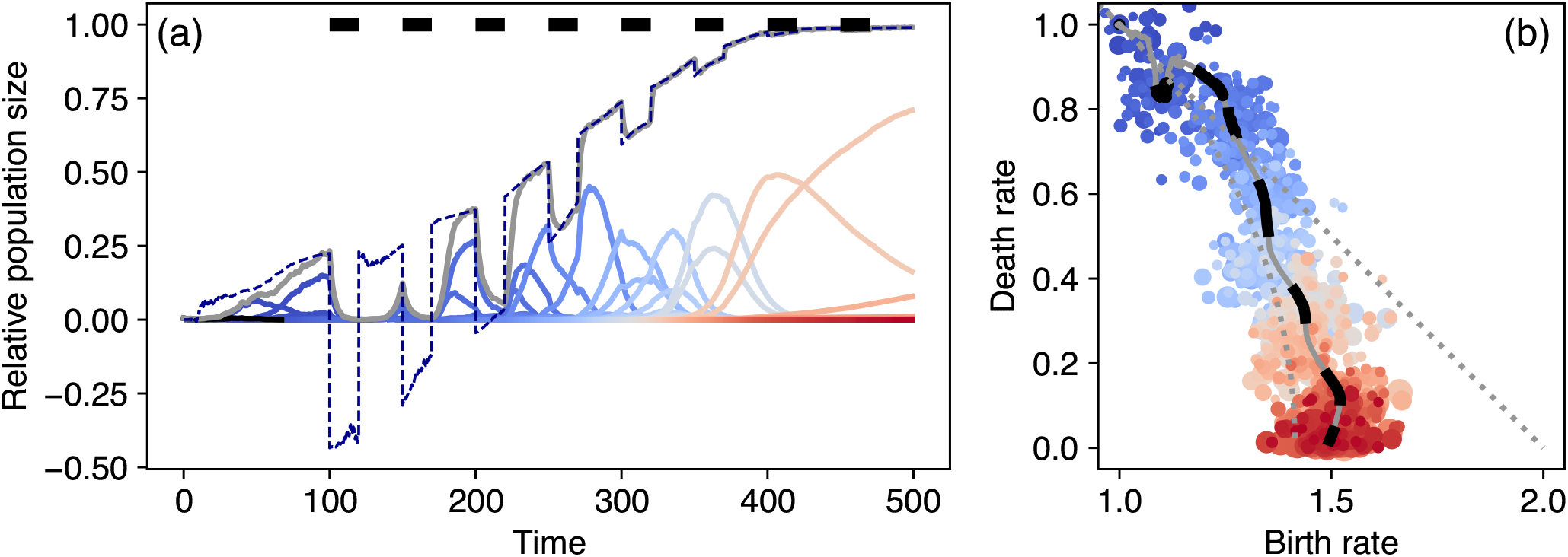
Exemplary dynamics for static treatment. Plot details and parameters as in Fig. 2. Black bars depict the times when Δ_*β*_ = 0.5. During treatment the effective carrying capacity can reduce to negative values. The population sizes, however, must be non-negative and thus approach zero when the effective carrying capacity becomes negative.

**Figure S6.**
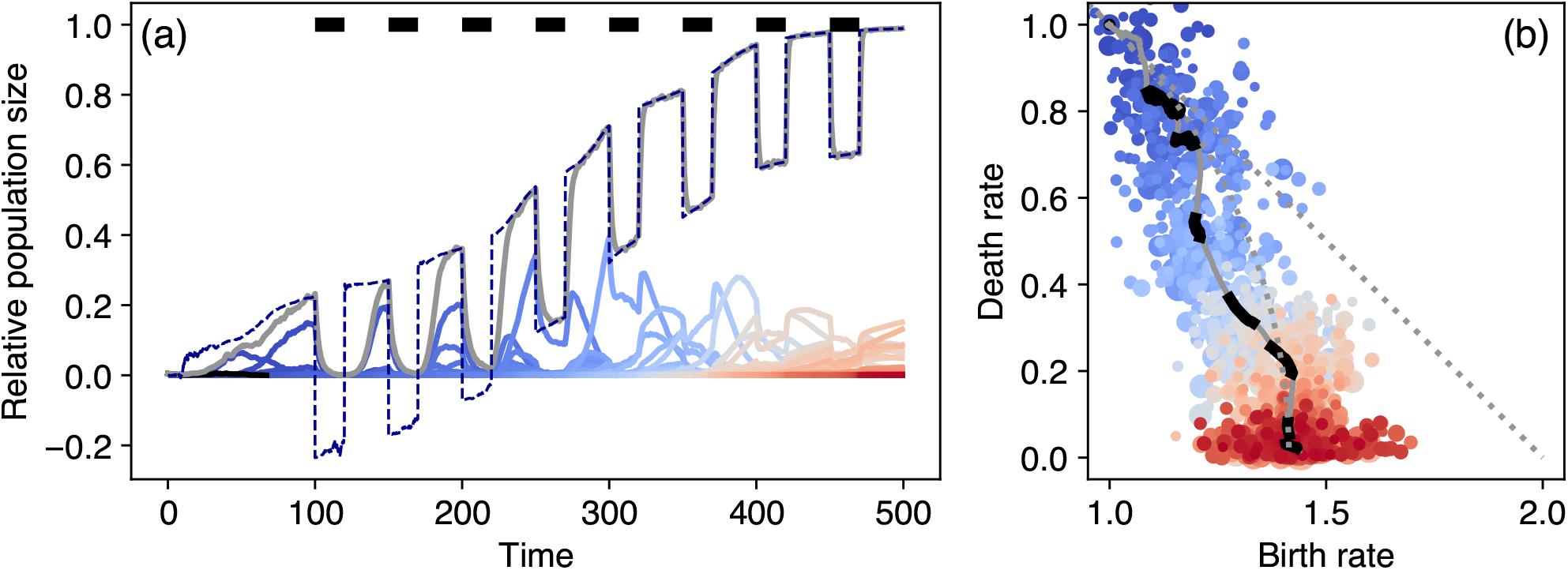
Exemplary dynamics for toxic treatment. Plot details and parameters as in Fig. 2. Black bars depict the times when Δ_*δ*_ = 0.5. During treatment the effective carrying capacity can reduce to negative values. The population sizes, however, must be non-negative values and thus approach zero when the effective carrying capacity becomes negative.

**Figure S7.**
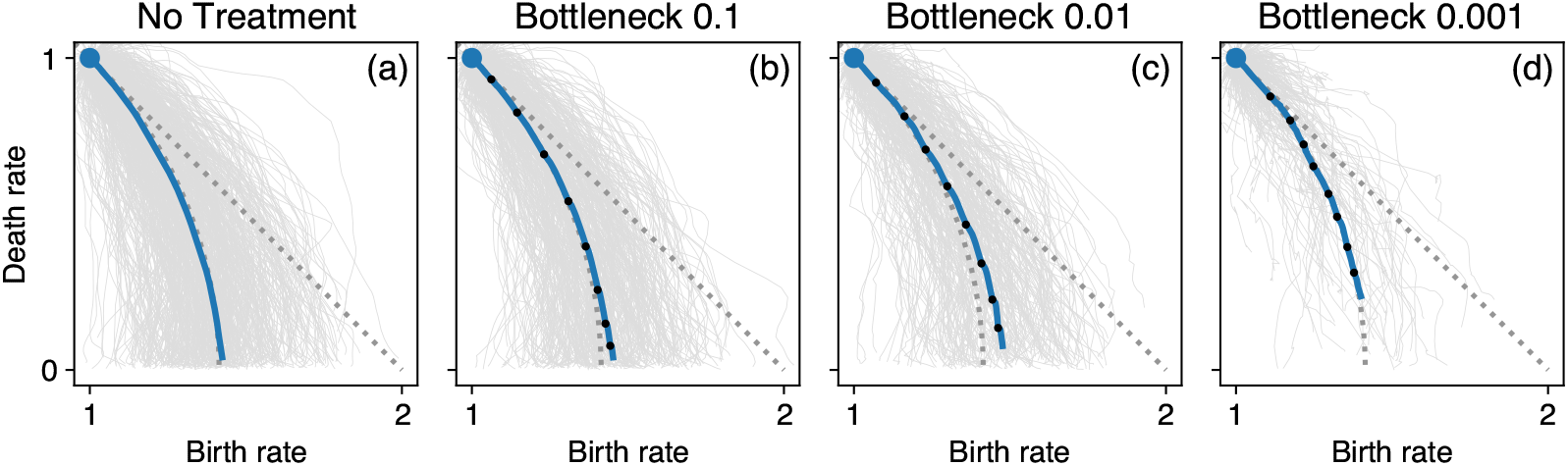
Trajectories of trait adaptation under density-affecting treatment. Grey lines represent the 1000 individual replicates. The thick lines show the ensemble average, blue stretches are treatment-off phases, black dots indicate the application of density-affecting treatment.

**Figure S8.**
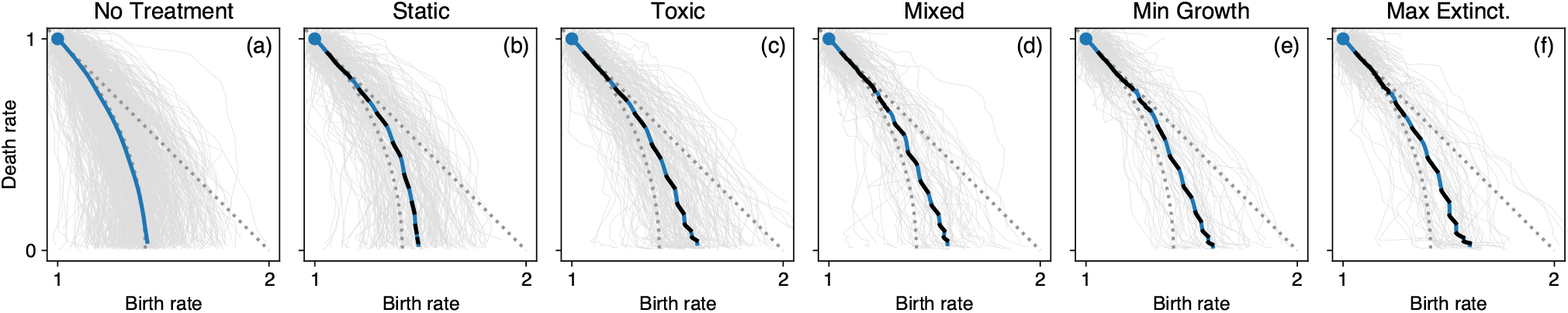
Trajectories of trait adaptation under trait-affecting treatment. Grey lines represent the 1000 individual replicates. The thick lines show the ensemble average, blue stretches are treatment-off phases, black stretches indicate treatment-on phases.

**Figure S9.**
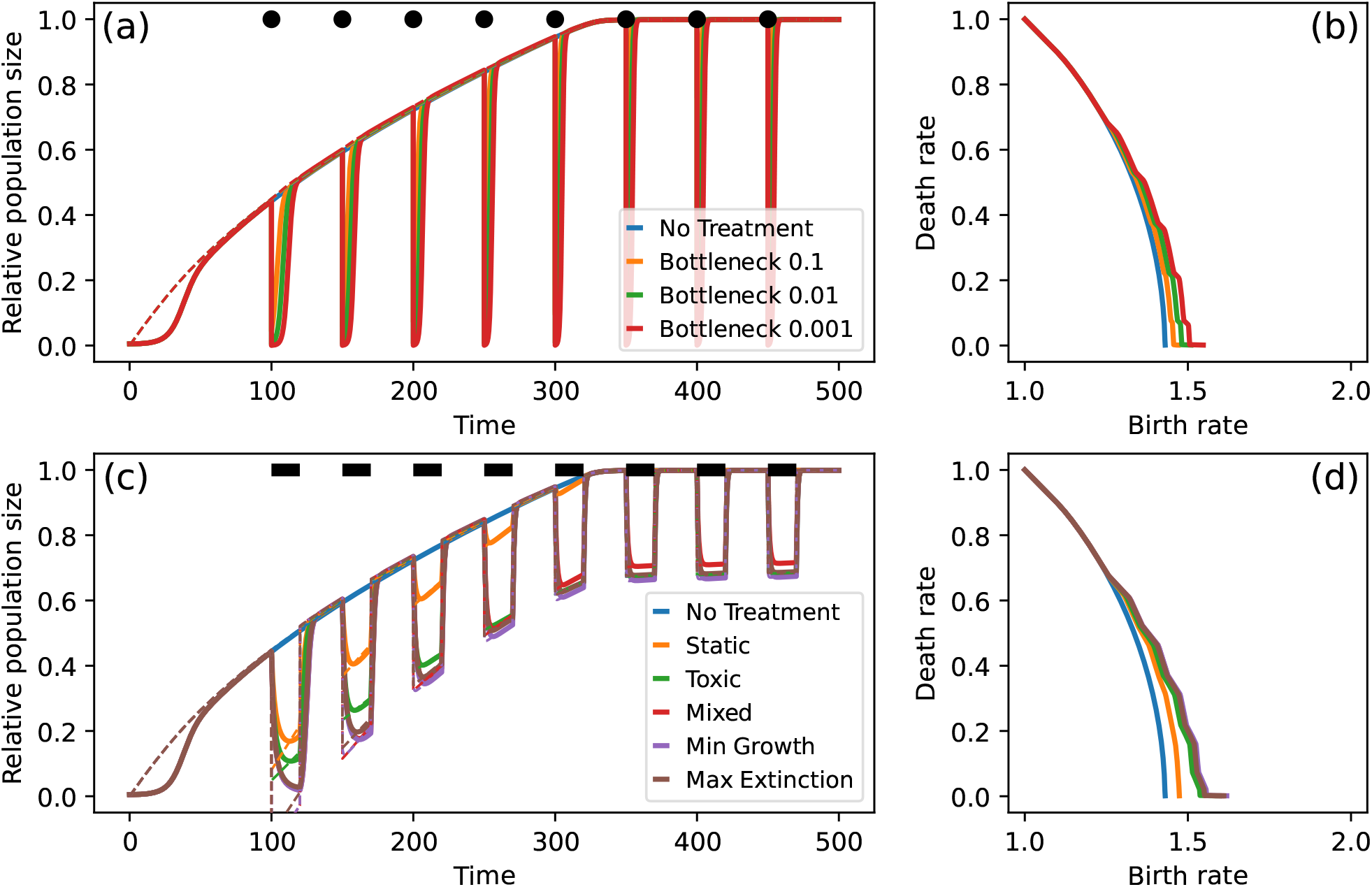
Deterministic adaptation dynamics under treatment - Net growth fitness gradient. Choosing the net growth gradient (Eq. (7)) as the fitness gradient in the deterministic model (Eq. 3) and parameter values from Tab. 1, we obtain adaptation dynamics that are similar to those presented for the stochastic model (Fig. 7).

**Figure S10.**
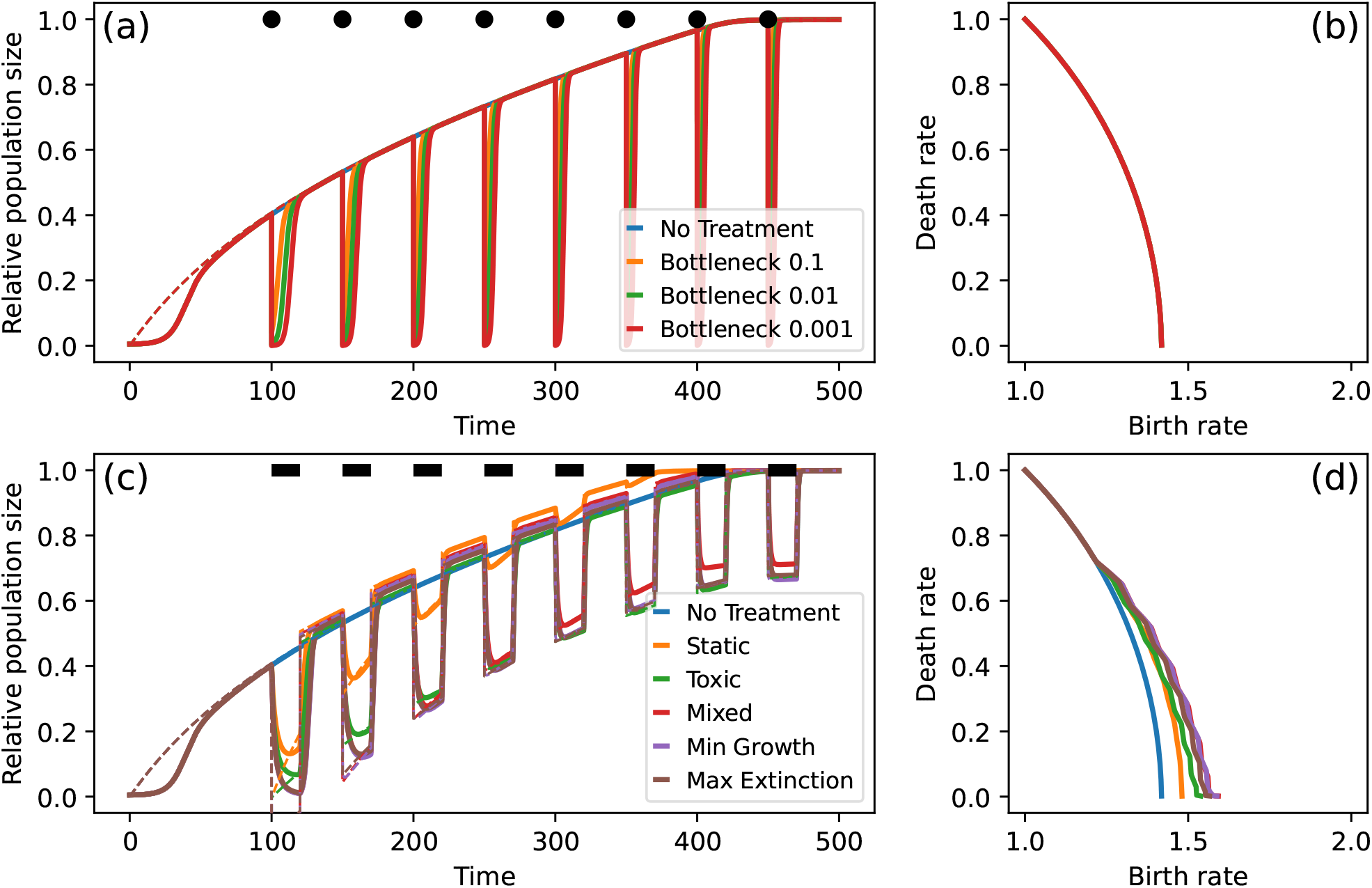
Deterministic adaptation dynamics under treatment - Survival probability fitness gradient. Choosing the survival probability gradient (Eq. (8)) as the fitness gradient in the deterministic model (Eq. 3) and parameter values from Tab. 1, we obtain adaptation dynamics that are similar to those presented for the stochastic model (Fig. 7). However, the density-affecting treatment type has no effect on the trait trajectory as the survival probability fitness gradient is density-independent.

**Figure S11.**
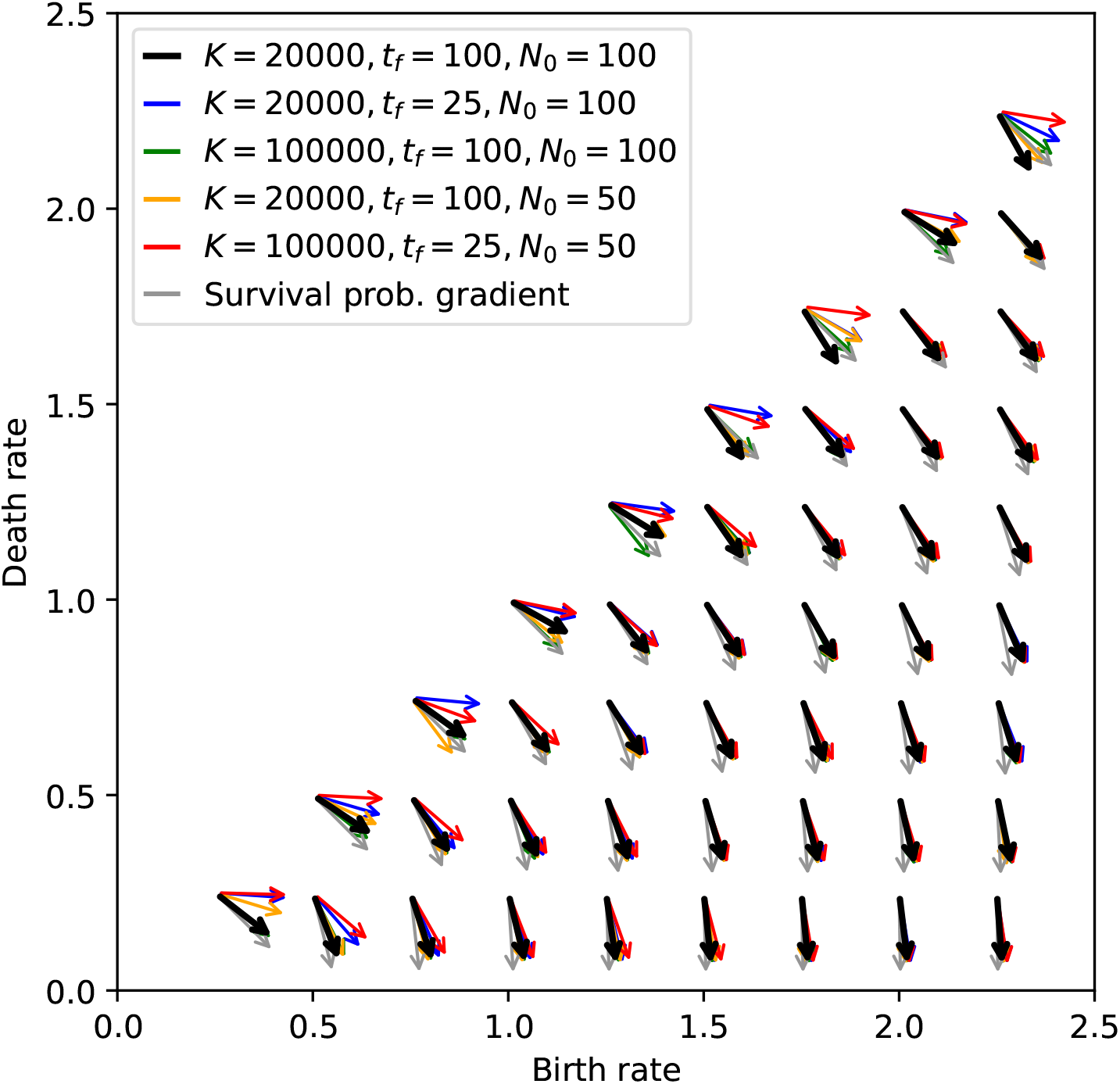
Observed initial steps of adaptation. Shown is the average direction of the adaptation trajectories in trait space until time *t*_*f*_ for different combinations of observation window *t*_*f*_, carrying capacity *K* and initial population size *N*_0_. Other parameters are chosen as given by Tab. 1. If the net growth was determining the adaptation trajectory, we expect adaptation steps that have a higher birth-rate component for decreasing density limitation (which can be realized by shorter observational window (blue arrows), higher carrying capacity (green arrows), smaller initial population size (yellow arrows) or all combined (red arrows)). If survival probability (grey arrows) was driving the adaptation we would expect the adaptation direction to not be affected by changes to *t*_*f*_, *K* or *N*_0_.

